# Cutting in-line with iron: ribosomal function and non-oxidative RNA cleavage

**DOI:** 10.1101/851097

**Authors:** Rebecca Guth-Metzler, Marcus S. Bray, Moran Frenkel-Pinter, Suttipong Suttapitugsakul, Claudia Montllor-Albalate, Jessica C. Bowman, Ronghu Wu, Amit R. Reddi, C. Denise Okafor, Jennifer B. Glass, Loren Dean Williams

## Abstract

Divalent metal cations are essential to the structure and function of the ribosome. Previous characterizations of the ribosome performed under standard laboratory conditions have implicated Mg^2+^ as a primary mediator of ribosomal structure and function. Possible contributions of Fe^2+^ as a ribosomal cofactor have been largely overlooked, despite the ribosome’s early evolution in a high Fe^2+^ environment, and its continued use by obligate anaerobes inhabiting high Fe^2+^ niches. Here we show that (i) Fe^2+^ cleaves RNA by in-line cleavage, a non-oxidative mechanism that has not previously been shown experimentally for this metal, (ii) the first-order rate constant with respect to divalent cations is more than 200 times greater with Fe^2+^ than with Mg^2+^, (iii) functional ribosomes are associated with Fe^2+^ after purification from cells grown under low O_2_ and high Fe^2+^, and (iv) a small fraction of Fe^2+^ that is associated with the ribosome is not exchangeable with surrounding divalent cations, presumably because it is tightly coordinated by rRNA and buried in the ribosome. In total, these results expand the ancient role of iron in biochemistry and highlight a possible new mechanism of iron toxicity.

**Key Points:**

1. Fe^2+^ cleaves rRNA by a non-oxidative in-line cleavage mechanism that is more than 200 times faster than in-line cleavage with Mg^2+^;
2. ribosomes purified from cells grown under low O_2_ and high Fe^2+^ retain ~10 Fe^2+^ ions per ribosome and produce as much protein as low O_2_, high Mg^2+^-grown ribosomes;
3. a small fraction (~2%) of Fe^2+^ that is associated with the ribosome is not exchangeable.

## Introduction

The translation system is responsible for the synthesis of all coded proteins and contains life’s most conserved ribonucleic acids. The common core of the ribosome is universal to all life (1, 2) and has been essentially invariant since the last universal common ancestor (3–5). Thus, ribosomes can be interrogated as molecular fossils (6–8). Because ribosomal structure and function are strongly dependent on divalent cations (M^2+^) (9), and because ribosomes originated long before the Great Oxidation Event (GOE), understanding ribosomal origins and evolution requires characterization of ribosomal interactions with M^2+^ ions under pre-GOE conditions (10–14).

In extant aerobic life, Mg^2+^ appears to be the dominant M^2+^ ion in the translation system. Hundreds of Mg^2+^ ions mediate ribosomal RNA (rRNA) folding and ribosomal assembly, in some instances binding to specific sites in the universal rRNA common core by direct coordination (9, 15–17). Mg^2+^ ions facilitate association of the large ribosomal subunit (LSU) and small ribosomal subunit (SSU) (18), stabilize folded tRNA (19), maintain the reading frame during translation (20), and link ribosomal proteins (rProteins) to rRNA (21). Mg^2+^ also catalyzes in-line cleavage (22–25), the reaction in which a divalent or trivalent metal catalyzes self-cleavage of RNA (26–29).

Before the GOE, anoxia would have stabilized abundant Fe^2+^ in the biosphere and hydrosphere. Under pre-GOE conditions, Fe^2+^ would not have caused the damage to biomolecules that occurs today in the presence of O_2_, via Fenton chemistry (30). In Fenton chemistry H_2_O_2_, a product of Fe^2+^ autoxidation by O_2_ (31), is reduced by Fe^2+^ generating hydroxyl radicals which can oxidatively damage nucleic acids (32–36). We recently reported that Fe^2+^ can fold RNA and mediate *in vitro* translation under “pre-GOE” conditions: in the presence of abundant Fe^2+^ and in the absence of O_2_ (37). Based on these findings, we proposed that early ribosomal folding and catalysis used Fe^2+^ instead of, or in combination with Mg^2+^ and other M^2+^ ions. However, we found lower translation rates with anoxic Fe^2+^ than with Mg^2+^ (37). While this observation could be partially explained by ribosomes adapting to the near-absence of Fe^2+^ and the presence of tens of mM Mg^2+^ in the ocean over the last two billion years, we assume that this explanation alone is insufficient because the catalytic core of the ribosome, and its central structure appears to be so conserved that it reflects the behavior of its evolutionary ancestor more than three billion years ago. One possible explanation for this finding is that non-oxidative damage of RNA mediated by Fe^2+^ is faster than with Mg^2+^.

Here we demonstrate that Fe^2+^ can damage RNA by in-line cleavage, which is distinct from previously characterized oxidative processes. We discovered that this second, non-oxidative mechanism of Fe^2+^-mediated RNA damage by in-line cleavage can be more extensive in some conditions than oxidative damage. We show that anoxic Fe^2+^ is efficient in catalyzing in-line cleavage, cleaving rRNA far more rapidly and extensively than Mg^2+^. Given that the reaction is likely first-order with respect to the metal (38, 39), the reaction rate constant (M^−1^ s^−1^) appears to be over 200 times greater for rRNA with Fe^2+^ than with Mg^2+^. The in-line mechanism of cleavage by Fe^2+^ was validated here by a variety of methods including reaction product characterization.

While metals such as Mg^2+^ cleave RNA, they are nonetheless essential for folding and function. In parallel with our experiments comparing Fe^2+^ and Mg^2+^ in-line cleavage, we investigated whether ribosomes from *E. coli* grown in pre-GOE conditions associate functionally with Fe^2+^ *in vivo*. We grew *E. coli* in anoxic conditions with ample Fe^2+^ in the growth media. We purified ribosomes from these bacteria and have probed their interactions with metals. We identified tightly bound M^2+^, which survive ribosomal purification. A small fraction (~2%) of Fe^2+^ ions are not exchangeable with Mg^2+^ in solution and are detectable after purification involving repeated washes in high [Mg^2+^] buffers. We use these tightly bound ions as reporters for more general M^2+^ association *in vivo*. The data are consistent with a model in which certain M^2+^ ions are deeply buried and highly coordinated within the ribosome (16). Our results suggest that ribosomes grown in pre-GOE conditions contain ~10 tightly bound Fe^2+^ ions compared to ~1 Fe^2+^ ion in ribosomes from standard growth conditions. Ribosomes washed with Fe^2+^ contained significantly higher Fe^2+^ and showed more rRNA degradation than ribosomes washed with Mg^2+^. Our combined results show the capacity for Fe^2+^ to (i) associate with functional ribosomes *in vivo* and *in vitro* and (ii) mediate significant non-oxidative damage. Our results have significant implications for the evolution of rRNA and iron toxicity in disease.

## Materials and Methods

### Ribosomal RNA purification

A one-tenth volume of sodium acetate (3.0 M, pH 5.2) and an equal volume of 25:24:1 phenol:chloroform:isoamyl alcohol, pH 5.2 (Fisher BioReagents) were added to purified ribosomes. The sample was vortexed and spun at 16,200 × g in a table-top centrifuge for 5 minutes. The top aqueous layer was transferred to a new tube and extracted twice in a 24:1 mixture of chloroform:isoamyl alcohol (Acros Organics) using the same procedure. rRNA was then precipitated by adding two volumes of 100% ethanol, followed by incubation at −20°C for 30 minutes. Precipitated rRNA was pelleted by centrifuging at 16,100 × g for 15 minutes. The pellet was washed with 70% ethanol and suspended in 0.1 mM sodium-EDTA (pH 8.0). Ribosomal RNA concentrations were quantified by A_260_ (1A_260_ = 40 μg rRNA mL^−1^).

### rRNA in-line cleavage reaction rates

Nuclease free water (IDT) was used in all experiments involving purified or transcribed RNA. rRNA for in-line cleavage experiments was purified as above by phenol-chloroform extraction followed by ethanol precipitation of commercial *E. coli* ribosomes (New England Biolabs). All in-line cleavage reaction solutions were prepared and incubated in the anoxic chamber. Fe and Mg solutions were prepared by dissolving a known mass of FeCl_2_-4H_2_O or MgCl_2_ salt in degassed water inside the chamber. 0.5 μg μL^−1^ of rRNA was suspended in degassed 20 mM HEPES pH 7.6, 30 mM KCl, 5% v/v glycerol [Invitrogen (UltraPure)], and either 25 mM of MgCl_2_ or 1 mM of FeCl_2_. Reactions were placed on a 37°C heat block and incubated for 4 days for the MgCl_2_ and no M^2+^ conditions and for 8 hours for the FeCl_2_ conditions. At each time point (0, 1.5, 3, 6, 12, 24, 48, and 96 hours for the MgCl_2_ and no M^2+^ conditions and 0, 7.5, 15, 30, 60, 120, 240, and 480 minutes for the FeCl_2_ conditions) 4.5 μL aliquots were combined with 0.5 μL of 1 M sodium phosphate buffer pH 7.6 to precipitate the Fe^2+^ or Mg^2+^ from solution and stored at −80°C. Aliquots were defrosted on ice and combined with 2X Gel Loading Buffer II (Amicon) then loaded onto a 1% Tris/Borate/EDTA agarose gel and run at 120V for 1.25 hours. The RNA in the gel was stained with GelStar™ (Lonza) and imaged with an Azure 6000 Imaging System (Azure Biosystems). Azurespot software was used as a pixel counter to create lane profiles. rRNA peaks were integrated by fitting to an Exponentially Modified Gaussian distribution using Igor Pro (v 7.08) **(Fig. S1)**. Observed pseudo first-order rate constants (k_obs_) were found by taking the negative of the slope from the natural logarithm of the normalized peak area vs. time plot. Reaction rate constants (k) were calculated by k = k_obs_/[M^2+^].

### In-line cleavage banding patterns

a-rRNA (40), which is composed of the core of the LSU rRNA, was synthesized and purified as previously described. Lyophilized a-rRNA was resuspended in degassed nuclease free water (IDT) inside the anoxic chamber. Fe and Mg solutions were prepared by dissolving known amounts of FeSO_4_-7H_2_O or MgSO_4_ in degassed nuclease free water inside the anoxic chamber. To initiate the reaction, 1 mM (final concentration) of Mg or Fe was added to 0.02 μg μL^−1^ a-rRNA in 20 mM HEPES-TRIS (pH 7.2) in a 37°C heat block. Samples were removed at 0, 0.25, 0.5, and 1 hr for added Fe^2+^, and at 24 hrs for added Mg^2+^. Divalent chelation beads (Hampton Research) were added to quench the reactions. Chelation beads were removed using spin columns. The RNA cleavage products were visualized using denaturing PAGE (6%, 8M urea) run at 120 V for ~1.3 hours stained with SYBR Green II.

### Fenton chemistry reactions

Purified rRNA from *E. coli* ribosomes (New England Biolabs) was obtained by phenol-chloroform extraction and ethanol precipitation as above. A stock solution of Fe/EDTA was prepared inside the anoxic chamber by dissolving a known amount of FeCl_2_-4H_2_O salt in degassed water then mixing with EDTA in degassed water. The Fe/EDTA was removed from the chamber for the Fenton reactions. Ribosomal RNA was suspended to 0.5 μg μL^−1^ in 20 mM HEPES pH 7.6, and 30 mM KCl, with 0% or 5% v/v glycerol and either 1 mM Fe/10 mM EDTA/10 mM ascorbate plus 0.3% v/v H_2_O_2_ or 10 mM EDTA as the reaction initiators wherein the initiators were separately dispensed onto the tube wall then vortexed with the other components. For the zero time points, reaction components were mixed in tubes containing the thiourea quenching agent at a final concentration of 100 mM. For non-zero time points the reaction mixtures were prepared as bulk solutions and incubated at 37°C on a heat block, after which aliquots were removed at 0, 10, and 60 minutes and mixed with the thiourea quenching agent at a final concentration of 100 mM. The stopped solutions were immediately frozen and stored at −80°C. For analysis, samples were defrosted on ice, combined with 2X Gel Loading Buffer II (Amicon), loaded onto a 1% Tris/Borate/EDTA agarose gel and run at 120V for 1.25 hours.

### Characterization of ApA cleavage products by HPLC

In-line cleavage reagents were prepared as previously described in the anoxic chamber. In duplicate reactions, 0.5 mM ApA RNA dinucleotide (5’ to 3’; TriLink BioTechnologies) was suspended in degassed 20 mM HEPES/NaOH pH 7.6, 30 mM KCl, 5% v/v glycerol, and combined with either water, 25 mM MgCl_2_ or FeCl_2_ (final concentration). Reactions were placed on a 37°C heat block with aliquots removed at 0, 0.25, 0.5, 1, 2, 4, and 8 days for no M^2+^ and Mg^2+^ samples and at 0, 0.25, 0.5, 1, and 2 days for Fe^2+^ samples. Aliquots were immediately quenched with 100 mM final concentration sodium phosphate pH 7.6, centrifuged at 2,000 × g for 1 minute, and the ApA-containing supernatant was collected to avoid transfer of Fe or Mg phosphate precipitate to the HPLC column. The samples were stored at −80°C prior to placement in the HPLC where they were held at 4°C. HPLC analyses were conducted on an Agilent 1260 Infinity HPLC with DAD UV-vis detector, with a path length of 1.0 cm. Products of the reactions were separated using a Kinetex XB-C18 column (150 × 2.1 mm, 2.6 μm particle size). The flow rate was 0.3 mL min^−1^ and the column temperature was held at 25°C. The mobile phase was water (0.1% formic acid) /acetonitrile. The gradient started with 100% water for the first 5 minutes and ramped to 55% acetonitrile over 25 minutes. The acetonitrile concentration was then ramped to 100% and was held as such for 10 minutes before returning to 100% water for column equilibration for 15 min. We recorded the elution at 210 nm, 220 nm, and 260 nm wavelengths, with a 180-400 nm spectrum detected in 2nm steps. To characterize reaction products, standards were spiked into product mixtures. The spiked standards were 0.5 mM ApA in 20 mM HEPES pH 7.6, 30 mM KCl, 5% v/v glycerol with either water, 2.5 μM adenosine, 3’-adenosine monophosphate, or 2’,3’-cyclic adenosine monophosphate.

### Characterization of ApA cleavage products by LC-MS

ApA was anoxically resuspended at 0.5 mM with 20 mM HEPES pH 7.6, 30 mM KCl, 5% v/v glycerol, and 25 mM FeCl_2_, and then incubated at 37°C for 2 days. The sample was analyzed by liquid chromatography mass spectrometry using and Agilent 1290 HPLC pump and thermostat; Agilent 1260 Autosampler and DAD UV-vis detector; path length: 0.6 cm; Agilent 1260 quaternary pump and RID; column: Phenomenex Kinetex 2.6 mmxB-C18100Å, LC column 150×2.1mm; column temp: 25°C; 10 μL injection with needle wash, 100 μL s^−1^ injection speed. The solvents were A) 0.1% formic acid in LC-MS grade water, and B) LC-MS grade acetonitrile a flow rates of 0.3 mL min^−1^; gradient: 5 min 100% A, 0% B; 20 min ramp to 45% A, 55% B; 10 min 0% A, 100% B; 1 min ramp 100% A, 0% B; 9 min 100% A, 0% B. Elutions were recorded at 210, 220 and 260 nm, with the entire spectrum (180-400 nm) detected in 2 nm steps. This system was coupled to an Agilent 6130 single quad MS Electrospray Ionization Mass Spectrometry system with scanning of ±65 to ±2000 m/z and capillary voltage of 2.0kV.

### Cell culture and harvesting

Culturing media consisted of LB broth (10 g L^−1^ NaCl, 10 g L^−1^ tryptone, 5 g L^−1^ yeast extract) amended with 4 mM tricine, 50 mM sodium fumarate, and 80 mM 3-(N-morpholino)propanesulfonic acid (MOPS; titrated with NaOH to pH 7.8). Fifty mL cultures containing all of these ingredients plus 0.25% v/v glycerol were inoculated from glycerol stocks of *Escherichia coli* MRE600 cells and shaken overnight at 37°C with or without O_2_ and with either 1 mM FeCl_2_ or ambient Fe^2+^ [6-9 μM, measured by the ferrozine assay (41)]. Two mL of each overnight culture was used to inoculate 1-L cultures in the same conditions. These cultures were then orbitally shaken at 37°C to OD_600_ 0.6-0.7. Aerobic cultures were grown in foil-covered Erlenmeyer flasks. Anaerobic fumarate-respiring cultures were inoculated into stoppered glass bottles containing medium that had been degassed with N2 for one hour to remove O_2_. Cells were then harvested by centrifugation at 4,415 × g for 10 minutes, washed in 20 mL buffer containing 10 mM Tris pH 7.4, 30 mM NaCl, and 1 mM EDTA, and pelleted at 10,000 × g for 10 minutes. Cell pellets were stored at −80°C until ribosome purification.

### Ribosome purification

The ribosome purification procedure was modified from Maguire et. al (42). All purification steps were performed in a Coy anoxic chamber (97% Ar, 3% H_2_ headspace) unless otherwise noted. Buffers varied in their metal cation content. The typical wash buffer contained 100 mM NH_4_Cl, 0.5 mM EDTA, 3 mM β-mercaptoethanol, 20 mM Tris pH 7.5, 3 mM MgCl_2_, and 22 mM NaCl. For “Fe purification” experiments, buffer was composed of 100 mM NH_4_Cl, 0.5 mM EDTA, 3 mM β-mercaptoethanol, 20 mM Tris pH 7.5, 1 mM FeCl_2_ and 28 mM NaCl. Sodium chloride concentrations were increased here to maintain the ionic strength of the buffer (131 mM). Elution buffers contained the same composition as the wash buffer except for NH_4_Cl (300 mM). Frozen cell pellets were resuspended in ribosome wash buffer and lysed in a BeadBug microtube compact homogenizer using 0.5 mm diameter zirconium beads (Benchmark Scientific). Cell lysate was transferred into centrifuge bottles inside the anoxic chamber which were tightly sealed to prevent O_2_ contamination. Cell debris were removed by centrifuging outside of the anoxic chamber at 30,000 × g for 30 minutes at 4°C. The soluble lysate was then transferred back into the chamber and loaded onto a column containing pre-equilibrated, cysteine-linked, SulfoLink™ Coupling Resin (Thermo Fisher Scientific). The resin was washed with 10 column volumes of wash buffer. Ribosomes were eluted into three 10 mL fractions with elution buffer. Eluted fractions were pooled inside the anoxic chamber into ultracentrifuge bottles which were tightly sealed. Ribosomes were pelleted outside the chamber by centrifuging at 302,000 × g for 3 hours at 4°C under vacuum in a Beckman Optima XPN-100 Ultracentrifuge using a Type 70 Ti rotor. Tubes containing ribosome pellets were brought back into the chamber and suspended in buffer containing 20 mM N-(2-hydroxyethyl)piperazine-N’-2-ethanesulfonic acid (HEPES; pH 7.6), 30 mM KCl, and 7 mM β-mercaptoethanol, heat-sealed in mylar bags, and stored at −80°C. Ribosome concentrations were calculated with a NanoDrop spectrophotometer assuming 1A_260_ = 60 μg ribosome mL^−1^ (conversion factor provided by New England Biolabs). This conversion factor was used to estimate the molecular mass of bacterial ribosomes, from which molarity was calculated. Biological triplicates of each growth and purification method were taken for downstream analyses.

### Ribosomal Fe content

Purified ribosomes were analyzed for iron content by total reflection X-ray fluorescence spectroscopy (TRXF) as described in Bray and Lenz et al (37).

### rProtein electrophoresis

For SDS-PAGE, purified ribosomes were normalized to 3.33 mg mL^−1^ in 2X SDS-PAGE dye, heated at 95°C for 5 minutes, and then incubated on ice for 2 minutes. Samples were loaded onto a 12% SDS acrylamide gel with a 4% stacking gel and run at 180 V for 60 minutes.

### In vitro translation

Translation reactions were based on the methods of Bray and Lenz et al. (37) with minor modifications. All 15 μL reactions contained 2.25 μL of purified ribosome samples normalized to 9 μg μL^−1^ (so that the final concentration of ribosomes in our reactions was 1.35 μg μL^−1^), 0.1 mM amino acid mix, 0.2 mM tRNAs, ~0.2 μg μL^−1^ of dihydrofolate reductase mRNA, and 3 μL of factor mix (with RNA polymerase, and transcription/translation factors in 10 mM Mg^2+^) from the PURExpress^®^ Δ Ribosome Kit (New England Biolabs). The reaction buffer was based on Shimizu et al. (43), with HEPES instead of phosphate buffer to avoid precipitation of metal phosphates. Buffer consisted of 20 mM HEPES (pH 7.3), 95 mM potassium glutamate, 5 mM NH_4_Cl, 0.5 mM CaCl_2_, 1 mM spermidine, 8 mM putrescine, 1 mM dithiothreitol (DTT), 2 mM adenosine triphosphate (ATP), 2 mM guanosine triphosphate (GTP), 1 mM uridine triphosphate (UTP), 1 mM cytidine triphosphate (CTP), 10 mM creatine phosphate (CP), and 53 μM 10-formyltetrahydrofolate. Divalent cation salts (MgCl_2_ or FeCl_2_) were added to 9 mM final concentration. The reaction buffer was lyophilized and stored at −80°C until resuspension in anoxic nuclease-free water immediately before experiments in the anoxic chamber. Reaction mixtures were assembled in the anoxic chamber and run at 37°C in a heat block for 120 minutes. Reactions were quenched on ice to terminate translation (43) and stored on ice until they were assayed for the extent of protein synthesis. Protein synthesis was measured using a DHFR assay kit (Sigma-Aldrich), which measures the oxidation of NADPH (60 mM) to NADP^+^ by dihydrofolic acid (51 μM). Assays were performed by adding 5 μL of protein synthesis reaction to 995 μL of 1X assay buffer. The NADPH absorbance peak at 340 nm (Abs_340_) was measured in 15 s intervals over 2.5 minutes. The slope of the linear regression of Abs_340_ vs. time was used to estimate protein activity (Abs_340_ min^−1^).

### Protein characterization by LC-MS/MS

After ribosomal purification, samples were reduced with β-mercaptoethanol, and then alkylated with 14 mM iodoacetamide (HEPES, pH 7.6) for 30 minutes at room temperature in the dark. Alkylation was quenched with 5 mM dithiothreitol for 15 minutes at room temperature in the dark. Proteins were purified by the methanol/chloroform precipitation method and were then digested with trypsin in a buffer containing 5% acetonitrile, 1.6 M urea, and 50 mM HEPES pH 8.8 at 37°C with shaking overnight. The digestion was quenched with addition of trifluoroacetic acid to a final concentration of ~0.2%. Peptides were purified by Stage-Tip (44) prior to LC-MS/MS analysis.

Peptides were dissolved in a solution containing 5% acetonitrile and 4% formic acid and loaded onto a C18-packed microcapillary column (Magic C18AQ, 3 μm, 200 Å, 75 μm × 16 cm, Michrom Bioresources) by a Dionex WPS-3000TPL RS autosampler (Thermostatted Pulled Loop Rapid Separation Nano/Capillary Autosampler). Peptides were separated by a Dionex UltiMate 3000 UHPLC system (Thermo Scientific) using a 112-minute gradient of 4-17% acetonitrile containing 0.125% formic acid. The LC was coupled to an LTQ Orbitrap Elite Hybrid Mass Spectrometer (Thermo Scientific) with Xcalibur software (version 3.0.63). MS analysis was performed with the data dependent Top15 method; for each cycle, a full MS scan with 60,000 resolution and 1 ×10^6^ AGC (automatic gain control) target in the Orbitrap cell was followed by up to 15 MS/MS scans in the Orbitrap cell for the most intense ions. Selected ions were excluded from further sequencing for 90 seconds. Ions with single or unassigned charge were not sequenced. Maximum ion accumulation time was 1,000 ms for each full MS scan, and 50 ms for each MS/MS scan.

Raw MS files were analyzed by MaxQuant (version 1.6.2.3; 45). MS spectra were searched against the *E. coli* database from UniProt containing common contaminants using the integrated Andromeda search engine (46). Due to the unavailability of the proteome database for *E. coli* strain MRE-600, the database for strain K12 was used. It has been shown that the two strains have nearly identical ribosome associated proteins (47). All samples were searched separately and set as individual experiments. Default parameters in MaxQuant were used, except the maximum number of missed cleavages was set at 3. Label-free quantification was enabled with the LFQ minimum ratio count of 1. The match-between-runs option was enabled. The false discovery rates (FDR) were kept at 0.01 at both the peptide and protein levels.

The results were processed using Perseus software (48). In the final dataset, the reverse hits and contaminants were removed. The LFQ intensity of each protein from the proteinGroups table was extracted and reported. For the volcano plots showing differential regulation of proteins, the ratios used were from the LFQ intensities of samples from each of the three experiments. The cutoff for differential expression was set at 2-fold. P-values were calculated using a two-sided T-test on biological triplicate measurements with the threshold p-value of 0.05 for significant regulation. The raw files are publicly available at http://www.peptideatlas.org/PASS/PASS01418 (username: PASS01418 and password: ZW2939nnw).

## Results

### In-line cleavage of rRNA: Mg^2+^ and anoxic Fe^2+^

By manipulating reaction conditions, we could switch the mode of rRNA cleavage between Fenton and in-line mechanisms. In-line is the only possible mechanism of cleavage by Mg^2+^ due to its fixed oxidation state and inability to generate hydroxyl radicals. We confirm the expectation that Mg^2+^-mediated in-line cleavage reactions are not inhibited by anoxia or hydroxyl radical quenchers (**Fig. S2**).

We confirm here in a variety of experiments that RNA is degraded by in-line cleavage when incubated with Fe^2+^ under anoxic conditions (**Fig. 1a**). Most of the experiments employed the rRNA of *E. coli* as substrate. A shorter RNA [a-RNA (40)] showed on a higher size resolution gel that RNA banding patterns and reaction products were nearly identical for Mg^2+^ and anoxic Fe^2+^ reactions (**Fig. 2**), indicating that preferred sites of cleavage are the same for both metals. Common sites of cleavage are indications of common mechanisms of cleavage (23). Neither Mg^2+^ nor anoxic Fe^2+^ cleavage was inhibited by glycerol (5%), which is known to quench hydroxyl radical and to inhibit hydroxyl radical cleavage (49). By contrast, glycerol inhibited cleavage by Fe^2+^ under conditions that favor Fenton-type cleavage **(Fig. S3)**. Glycerol did not inhibit Mg^2+^ in-line cleavage under any conditions **(Fig. S2)**.

**Figure 1.**
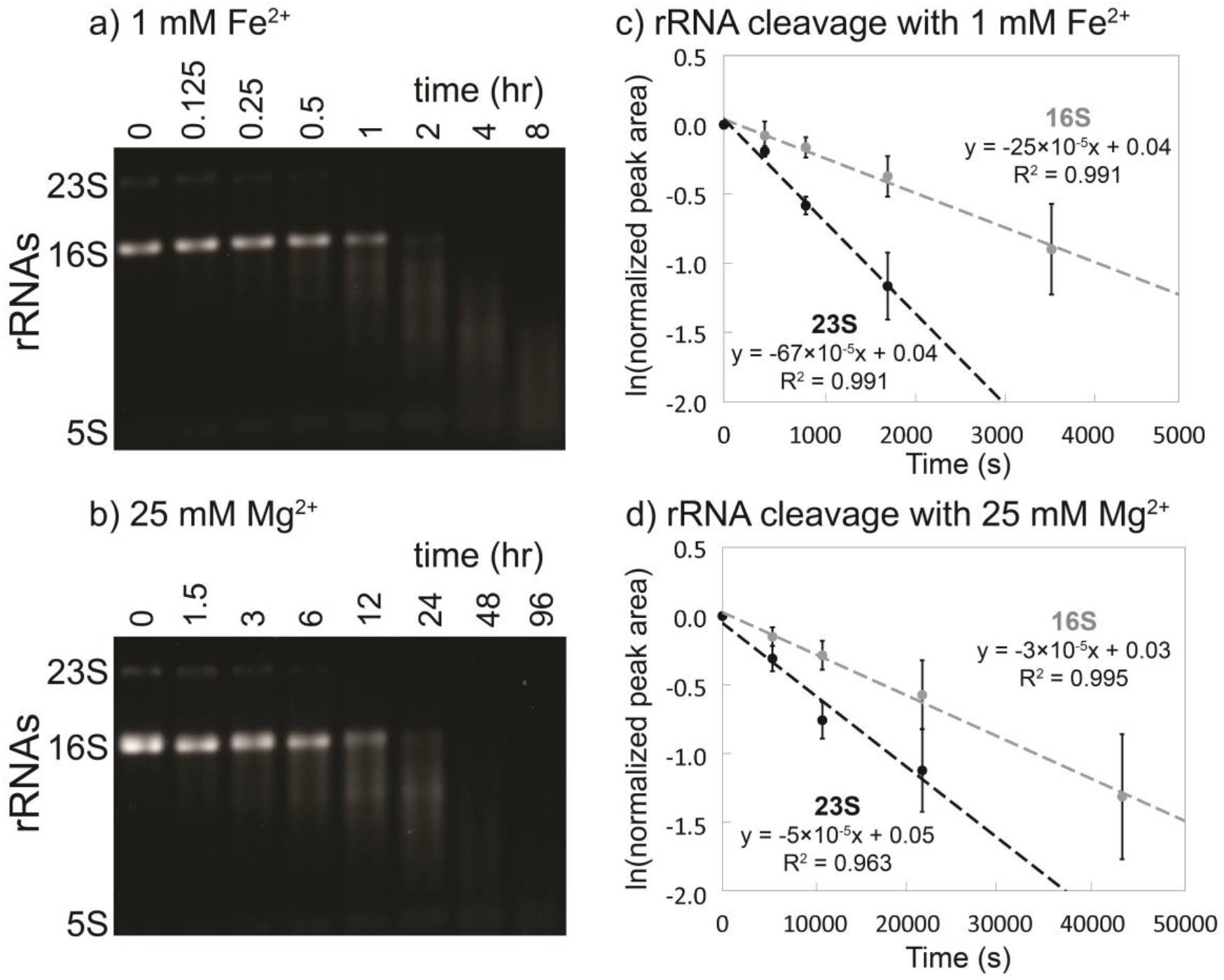
In-line cleavage of rRNA in anoxia. In-line cleavage of purified rRNAs with a) 1 mM Fe^2+^ (0-8 hr) and b) 25 mM Mg^2+^ (0-96 hr). Reactions were conducted in an anoxic chamber at 37°C in the presence of the hydroxyl radical quencher glycerol (5% v/v) and were analyzed by 1% agarose gels. Pseudo first-order rate plots were extracted from 23S and 16S band intensity for c) 1 mM Fe^2+^ and d) 25 mM Mg^2+^ conditions. The Mg^2+^ time axis is 10 times greater than the Fe^2+^ time axis. Error bars represent the S.E.M. (n = 3).

**Figure 2.**
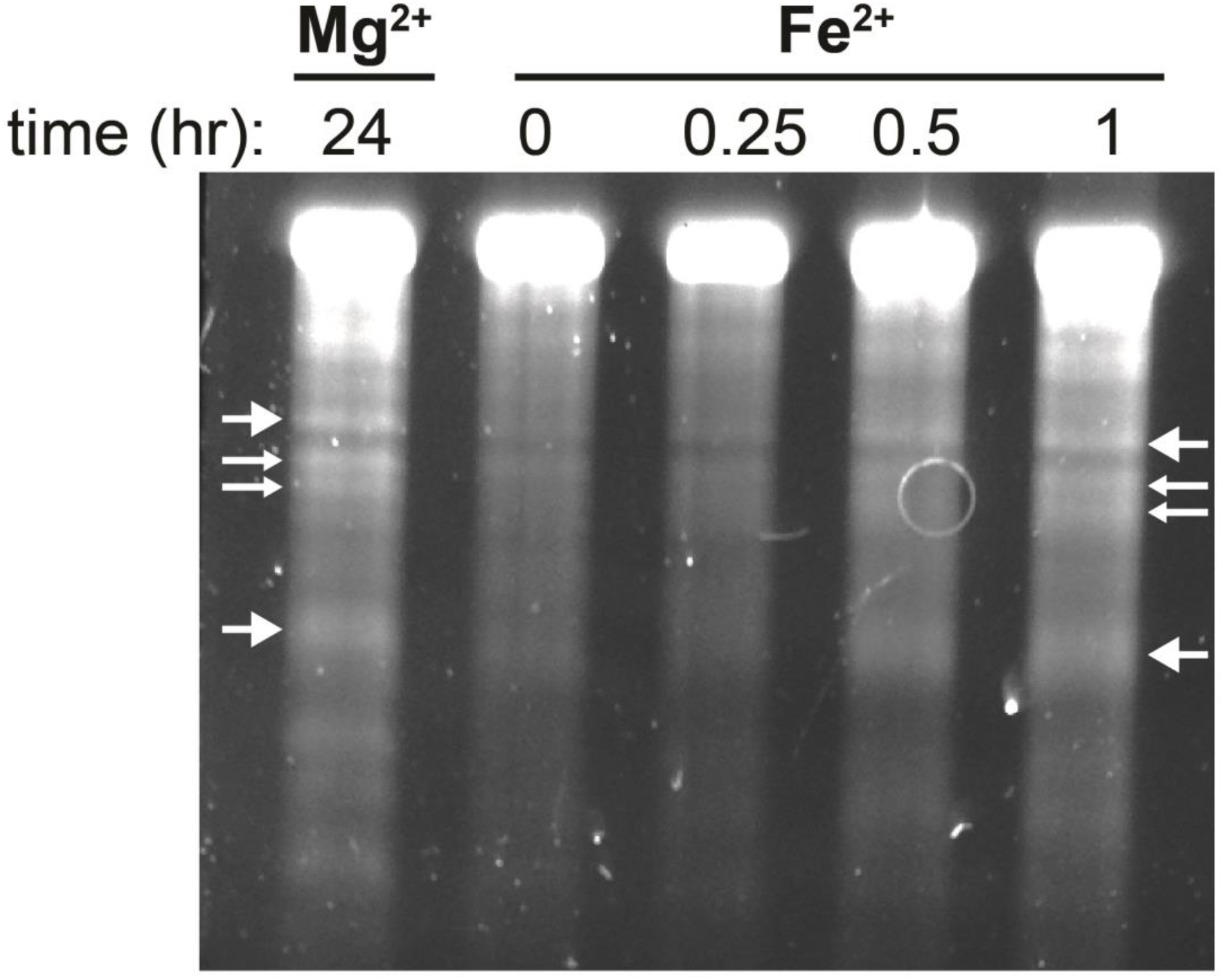
In-line cleavage banding patterns are the same for rRNA cleavage with Mg^2+^ and anoxic Fe^2+^. Several primary cleavage bands of a-rRNA (40) are indicated by arrows. This gel is 6% polyacrylamide, 8 M urea showing in-line cleavage mediated by 1 mM Mg^2+^ or 1 mM anoxic Fe^2+^ at 37°C for varying amounts of time. Reactions were run in 20 mM Tris-HEPES, pH 7.2.

In the absence of O_2_, cleavage rates are significantly greater for Fe^2+^ than for Mg^2+^. For 16S and 23S rRNAs, 1 mM Fe^2+^ caused significant in-line cleavage of rRNA after 30 minutes at 37°C. Both rRNAs were completely degraded after 2 hours in anoxic Fe^2+^ (**Fig. 1a**). By contrast, when the M^2+^ ion was switched from 1 mM Fe^2+^ to 25 mM Mg^2+^, only a modest amount of in-line cleavage was observed after 6 hours (**Fig. 1b**). Fitting of the data to a pseudo first-order rate model **(Fig. 1c and 1d)** reveals apparent rate constants for cleavage of the full-length 23S rRNA with Fe^2+^ is 67 × 10^−5^ s^−1^ and with Mg^2+^ is 5 × 10^−5^ s^−1^. The rate constants for cleavage of the 16S rRNA is 25 × 10^−5^ s^−1^ for Fe^2+^ and 3 × 10^−5^ s^−1^ for Mg^2+^ (**Table 1**). These apparent rate constants do not account for differences in metal concentration or in RNA length.

**Table 1:** rRNA cleavage pseudo first-order rate constants.

| Metal | rRNA | $k_{\text{obs}}$ ($10^{-5} \text{ s}^{-1}$) | S.E.M. ($10^{-5} \text{ s}^{-1}$) |
| --- | --- | --- | --- |
| 1 mM $\text{Fe}^{2+}$ | | | |
|  | 23S | 67 | 12 |
|  | 16S | 25 | 8.2 |
| 25 mM $\text{Mg}^{2+}$ | | | |
|  | 23S | 5 | 1.1 |
|  | 16S | 3 | 0.8 |

In sum, reactions with Mg^2+^ and anoxic Fe^2+^ showed a lack of inhibition by a hydroxyl radical quencher. By contrast, the quencher inhibited reactions with Fe^2+^ in the presence of O_2_. The observed rate constant for in-line cleavage for rRNA is ~10 times greater for 1 mM Fe^2+^ than for 25 mM Mg^2+^. Under these conditions, in-line cleavage is expected to scale with metal concentration (38, 39) so that that the reaction rate constant, k (M^−1^ s^−1^), is increased with Fe^2+^ by ~300 times for the 23S and by ~200 times for the 16S. We demonstrate that although Fe^2+^ interacts in the same way as Mg^2+^ with RNA, the cleavage potency of Fe^2+^ is greatly enhanced.

### Characterization of in-line cleavage reaction products

To confirm that Fe^2+^ catalyzes non-oxidative in-line RNA cleavage, a series of cleavage reactions were performed on the dinucleotide ApA. The products of the reaction were characterized by HPLC via spiking with standards, and by LC-MS. A small RNA with only one possible cleavage site allowed us to identify specific cleavage products, which report on the mechanism of scission. In-line cleavage leads to 2’,3’-cyclic phosphate upstream of the scission site and a downstream 5’OH RNA fragment. Subsequently, the 2’,3’-cyclic phosphate can hydrolyze to either a 2’ or 3’ monophosphate. Conversely, oxidative cleavage of RNA by a hydroxyl radical that may be formed during iron oxidative processes abstracts a proton from a ribose sugar leading to a variety of products but not 2’,3’-cyclic phosphate, 2’-phosphate, or 3’-phosphate RNA fragment (50). Anoxic incubation of ApA with 25 mM Mg^2+^ or Fe^2+^ produces adenosine, 2’,3’-cyclic adenosine monophosphate (2’,3’-cAMP), and 3’-adenosine monophosphate (3’-AMP) (**Fig. 3a-d** and **Fig. S4**). The repertoire of products is matched for the two metals, pointing to a common in-line cleavage mechanism.

**Figure 3.**
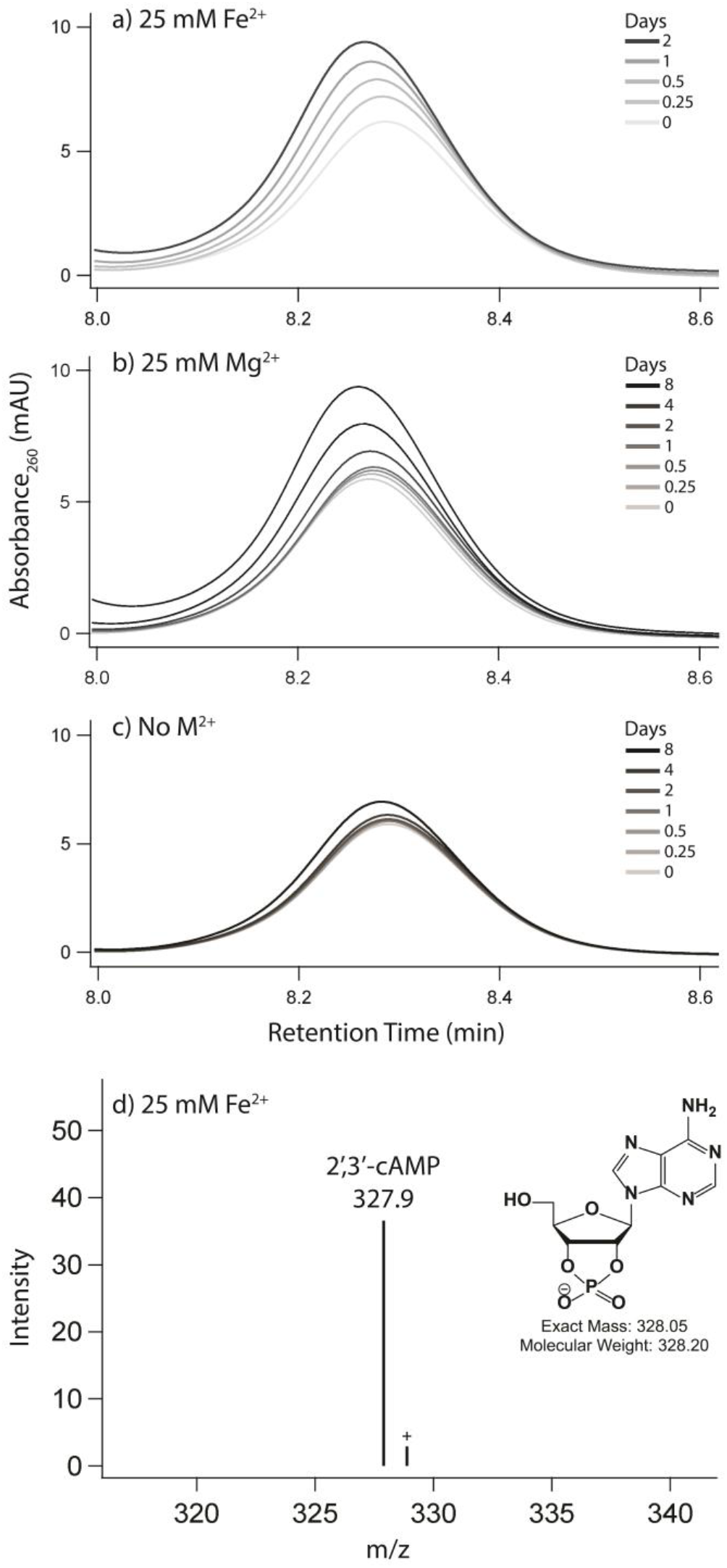
2’,3’-cAMP is formed upon incubation of ApA with Fe^2+^ or Mg^2+^. HPLC chromatograms show the accumulation of 2’,3’-cAMP, a direct product of an in-line cleavage mechanism, upon incubation of ApA with either a) 25 mM Fe^2+^, b) 25 mM Mg^2+^, or c) no metal for the negative control. Panel d shows identification of the 2’,3’-cyclic adenosine monophosphate by LC-MS of ApA incubated with 25 mM Fe^2+^ for 2 days. Labeled species correspond to [M-H]^−^ ions. Reactions were incubated anoxically at 37°C in the presence of 5% (v/v) glycerol.

### M^2+^ exchange during ribosomal purification

We hypothesized that O_2_ and Fe^2+^ content during bacterial growth could affect the iron content of ribosomes. However, the vast majority of ribosomal M^2+^ ions are exchangeable (51) and canonical ribosome purification procedures use high Mg^2+^ buffers (52) to maintain folding and stability. Therefore, spontaneous exchange of *in vivo* bound M^2+^ with those in the buffer occurs during purification, suggesting that the final Fe^2+^ content of purified ribosomes depends on the type of M^2+^ in the purification buffer. Indeed, ribosomes purified in solutions with 1 mM Fe^2+^ contained significantly higher Fe^2+^ than those purified in 3 mM Mg^2+^ regardless of growth condition (**Fig. 4**). All ribosome samples purified in 1 mM Fe^2+^ contained similar Fe^2+^ (~400-600 mol Fe mol^−1^ ribosome). These results show that the vast majority of ribosomal M^2+^ ions are exchangeable and that M^2+^ exchange takes place during purification.

**Figure 4.**
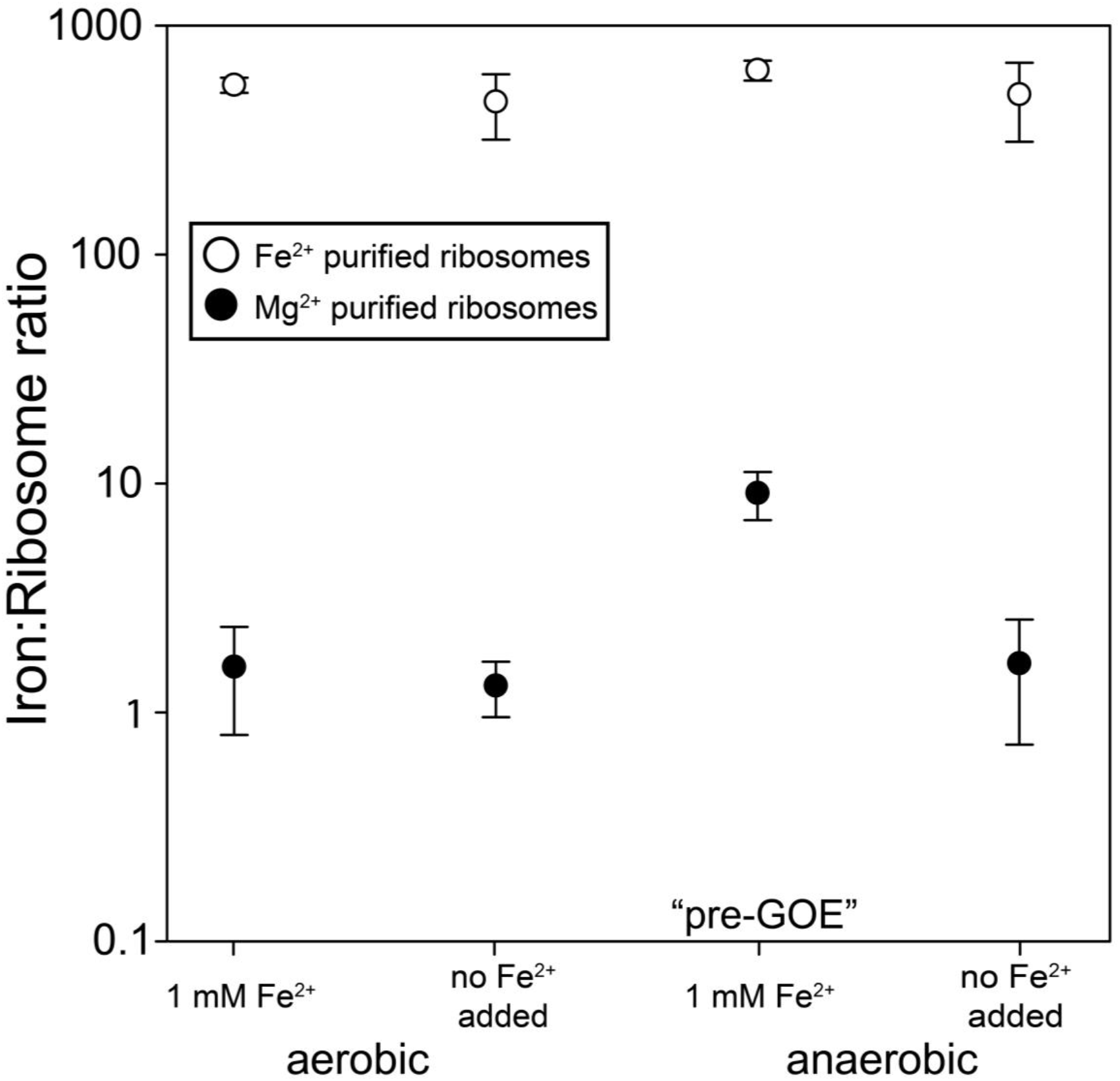
Iron content (mol Fe mol^−1^ ribosome) of purified ribosomes. *E. coli* were grown aerobically or anaerobically at 1 mM Fe^2+^ or ambient Fe^2+^ (6-9 μM, no Fe added), and purified in buffers containing either 3 mM Mg^2+^ (black circles) or 1 mM Fe^2+^ (white circles). Error bars represent the S.E.M. (n=3).

### Tight ribosomal binding of a subset of M^2+^

A small subset of ribosomal M^2+^ ions are not exchangeable during purification. Ribosomes retain this subset of *in vivo* divalent cations after purification. We harvested *E. coli* in log phase from four growth conditions: oxic or anoxic with high Fe^2+^ in the medium (1 mM Fe^2+^), and oxic or anoxic without added Fe^2+^ in the growth medium (6-9 μM Fe^2+^). Ribosomes from *E. coli* grown in pre-GOE conditions (anoxic, high Fe^2+^) contained quantitatively reproducible elevated levels of Fe^2+^ after purification in solutions containing Mg^2+^. We detected around 9 mol Fe mol^−1^ ribosome from cells grown in pre-GOE conditions purified in solutions with high Mg^2+^ (**Fig. 4)**. The three other growth conditions yielded ribosomes containing near background levels of Fe^2+^ (< 2 mol Fe mol^−1^ ribosome).

### Quantitating translation

Ribosomes from all four growth conditions produced active protein in translation assays. Ribosomes were functional *in vitro* under standard conditions (with 10 mM Mg^2+^) and also in 8 mM Fe^2+^ + 2 mM Mg^2+^ under anoxia. Regardless of whether translation activity was assayed in the presence of 10 mM Mg^2+^ or 8 mM Fe^2+^ + 2 mM Mg^2+^, ribosomes synthesized aerobically in the absence of Fe^2+^ have higher activity than the ribosomes synthesized anaerobically in the absence of Fe^2+^ (p< 0.08; **Fig. S5**). The presence of 1 mM Fe^2+^ in the bacterial growth conditions did not affect ribosomal activity; the only differences are whether growth conditions are aerobic or anaerobic, and whether 10 mM Mg^2+^ or 8 mM Fe^2+^ + 2 mM Mg^2+^ are used in the assay. Translation was reduced in the presence of Fe^2+^ compared to Mg^2+^, consistent with our previous work (37). The translational activity of ribosomes harvested from anaerobic cells was slightly less than from those from aerobic cells. Ribosomes from all four growth conditions contained intact 23S, 16S, and 5S rRNAs with purification in 3 mM Mg^2+^ (**Fig. 5a**) resulting in a higher proportion of intact rRNA relative to purification in 1 mM Fe^2+^ (**Fig. 5b**). Each purification also contained a full suite of rProteins as indicated by mass spectrometric analysis and by gel electrophoresis (**Fig. S6**). The protein composition of ribosomes from 1 mM Fe^2+^ growth conditions (**Fig. S6b**) was similar to that from Mg^2+^ growth conditions (**Fig. S6a**).

**Figure 5.**
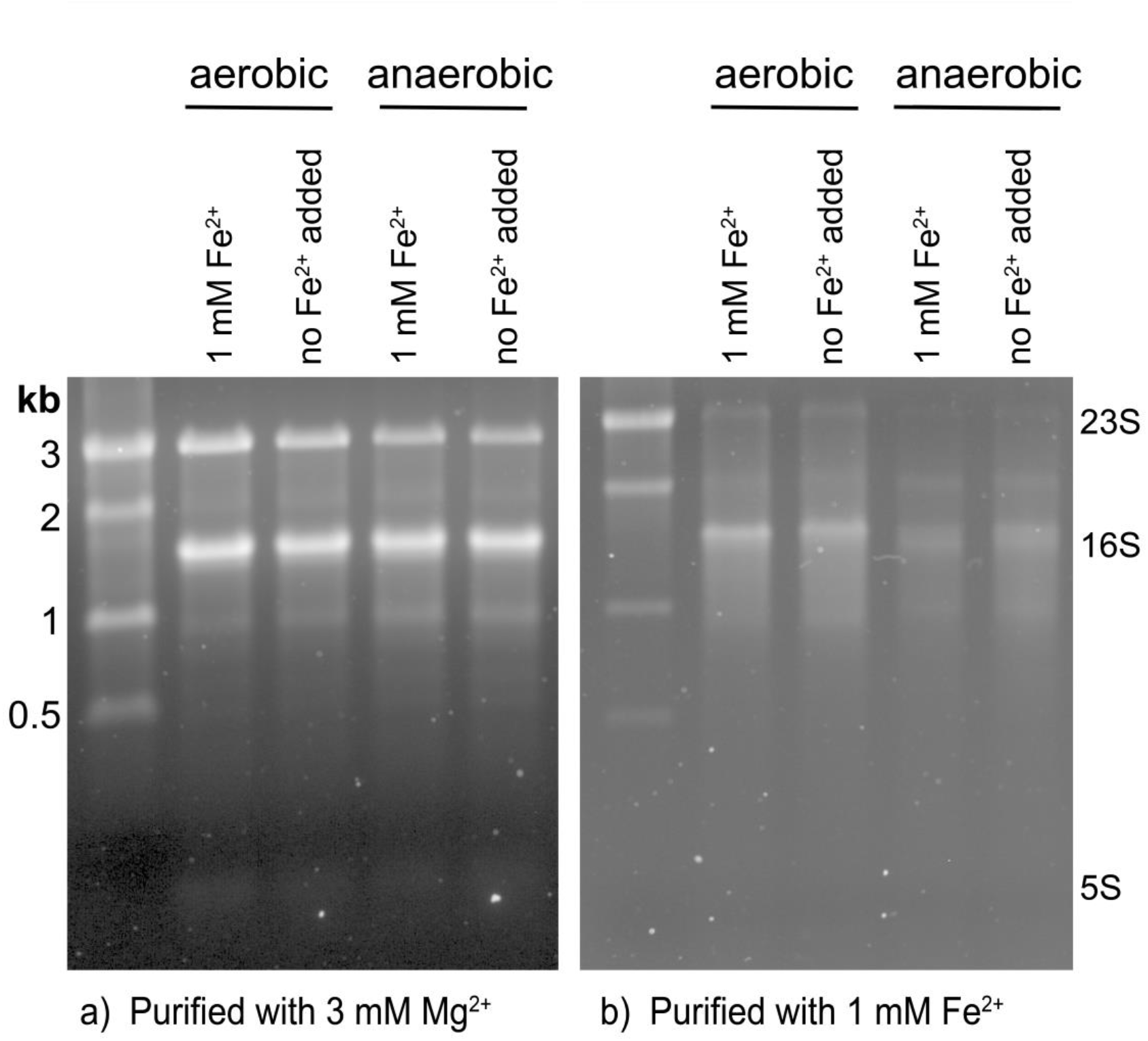
1% agarose gels showing rRNA from ribosomes purified in (a) 3 mM Mg^2+^ and (b) 1 mM Fe^2+^. The banding pattern suggests that rRNA is relatively more intact in ribosomes purified with 3 mM Mg^2+^ than in ribosomes purified with 1 mM Fe^2+^.

### rProtein characterization

In addition to oxidative mechanism, our results pointed to a non-oxidative cleavage mechanism of RNA with Fe^2+^. So, we next asked whether ribosomes might adopt different proteins to cope with high Fe^2+^ in both oxic and anoxic conditions. Ribosomes under all four growth conditions contained a full repertoire of rProteins, and were associated with additional proteins, as determined by mass spectrometry. These non-ribosomal proteins ranged in function from translation to central metabolism. Proteins from anaerobic pathways were generally more abundant in ribosomes from anaerobic cells while proteins from aerobic pathways were more abundant in ribosomes from aerobic cells (**Tables S1, S2**). Proteins for synthesis of enterobactin, an Fe^3+^-binding siderophore, were more abundant in ribosomes from aerobic cells and from those grown without the addition of Fe, while the bacterial non-heme ferritin subunit was more abundant in ribosomes from anaerobic cells regardless of the Fe^2+^ content in the media (**Table S2**). Several proteins were differentially expressed in ribosomes grown in pre-GOE conditions relative to other growth conditions (**Fig. S7**). Notably, ribosomes grown anaerobically with high Fe^2+^ had five times the abundance of the protein YceD than ribosomes grown anaerobically without added Fe^2+^. Anaerobic high Fe^2+^ ribosomes had one third the abundance of the rProtein S12 methylthiotransferase protein RimO and rRNA LSU methyltransferase K/L protein RlmL than ribosomes from aerobically grown cells with 1 mM Fe^2+^.

## Discussion

### Iron promotes rapid in-line cleavage of rRNA

Mg^2+^ is known to cleave the RNA phosphodiester backbone via an in-line mechanism (22, 23). We have shown here that Fe^2+^, like Mg^2+^, can cleave RNA by a non-oxidative in-line mechanism. We used cleavage of 23S and 16S rRNA to determine the observed rate constants of both Mg^2+^- and Fe^2+^-mediated cleavage. The k_obs_, uncorrected for metal concentration, for in-line cleavage by Fe^2+^ is around 10 times greater than for Mg^2+^. Previous studies of metal concentration effects on k_obs_ suggest that in-line cleavage is first-order with respect to metal concentration (38, 39), allowing the calculation of a per molar metal reaction rate constant by k = k_obs_/[M^2+^]. Assuming this first-order relationship in our experiments, k with Fe^2+^ is ~300 times greater for the 23S and ~200 times greater for the 16S than with Mg^2+^. In **Table S3** we compare our results to literature k values of Mg^2+^ and Zn^2+^ in-line cleavage taken under a range of conditions (38, 39, 53–57), normalizing for the number of cleavable phosphates in the RNA substrate. Changes in metal identity, RNA length, RNA folding, pH, and temperature, result in differences in normalized rate constants. The values extend over four orders of magnitude. Rate enhancement by switching Mg^2+^ to another metal while other conditions are held constant is greater for Fe^2+^ than for Zn^2+^, highlighting the rapidity of cleavage by Fe^2+^.

Support for a non-oxidative in-line mechanism of cleavage of RNA by anoxic Fe^2+^ is provided by observations that the rate of the reaction is not attenuated by anoxia and that the sites of cleavage appear to be conserved for Mg^2+^ and anoxic Fe^2+^. The absence of hydroxyl radical intermediates in the anoxic cleavage reaction is confirmed by the lack of inhibition by a hydroxyl radical quencher known to inhibit Fenton chemistry (32). Cleavage products of the RNA dinucleotide ApA include only those that are expected from an in-line mechanism and align with products formed with Mg^2+^. Among these is 2’,3’-cyclic phosphate, the hallmark of in-line attack of the bridging phosphate by the 2’OH.

In-line cleavage is the dominant mechanism of Fe^2+^ cleavage when contributions from Fenton-mediated processes are minimized and is the only mechanism of Mg^2+^ cleavage. By contrast, in oxic environments, transient Fe^2+^ oxidation generates hydroxyl radicals (31) that cleave nucleic acids (30, 32–35). Our results have significant implications for iron toxicity and human disease. The potency of Fe^2+^ in inducing rRNA cleavage may lead to decreased longevity of Fe^2+^-containing ribosomes. In fact, rRNA cleavage linked to Fe^2+^ oxidation, as in the human ribosome in Alzheimer’s disease (58), or in yeast rRNA (59), could be in some measure attributable to Fe^2+^ in-line cleavage.

Fe^2+^ appears to be a potent all-around cofactor for nucleic acids. The combined results indicate that:

a. rRNA folds at lower concentration of Fe^2+^ than Mg^2+^ (37),
b. at least a subset of ribozymes and DNAzymes are more active in Fe^2+^ than in Mg^2+^ (60, 61),
c. the translation system is functional when Fe^2+^ is the dominant divalent cation (37),
d. at low concentrations of M^2+^, T7 RNA polymerase is more active with Fe^2+^ than with Mg^2+^ (62),
e. a broad variety of nucleic acid processing enzymes are active with Fe^2+^ instead of Mg^2+^ (62),
f. rates of in-line cleavage are significantly greater for Fe^2+^ than for Mg^2+^ (here), and
g. Fe^2+^ but not Mg^2+^ confers oxidoreductase functionality to some RNAs (17, 63).

### Why so fast?

Our previous DFT computations (62) help explain why Fe^2+^ is such a potent cofactor for RNA. Conformations and geometries of coordination complexes with water and/or phosphate are nearly identical for Fe^2+^ or Mg^2+^. However, differences between Mg^2+^ and Fe^2+^ are seen in the electronic structures of coordination complexes.

Firstly, because of low lying d orbitals, Fe^2+^ has greater electron withdrawing power than Mg^2+^ from first shell phosphate ligands. In coordination complexes with phosphate groups, the phosphorus atom is a better electrophile when M^2+^ = Fe^2+^ than when M^2+^ = Mg^2+^. This difference between Mg^2+^ and Fe^2+^ is apparent in both ribozyme reactions and in-line cleavage reactions.

Secondly, Fe^2+^(H_2_O)_6_ is a stronger acid than Mg^2+^(H_2_O)_6_; depletion of electrons is greater from water molecules that coordinate Fe^2+^ than from those that coordinate Mg^2+^. The lower pKa of Fe^2+^(H_2_O)_6_ may promote protonation of the 5’OH leaving group during cleavage. Metal hydrates with low pKa’s have been reported to induce RNA cleavage better than less acidic metal hydrates (22).

In in-line cleavage, RNA coordinates M^2+^ or the M^2+^ hydrate (22, 23). Indeed, studies of the in-line cleavage fragment patterns have previously been used to probe structural information on RNA molecules, such as metal-binding sites (26, 27). We demonstrated with ApA that RNA secondary structure is not required for in-line cleavage. The same activities that drive in-line cleavage (e.g. 2’OH activation and coordination of the leaving group) are thought to occur in metal-catalyzed ribozyme cleavage (64). Multiple ribozymes (60) and DNAzymes (61) have been observed to function with Fe^2+^ as a cofactor. Our results with Fe^2+^ in-line cleavage, and in-line cleavage in general, require no enzymatic activity.

The remarkably high cleavage activity of Fe^2+^ with RNA demonstrated here bears relevance to prebiotic chemistry and early biochemistry. Because these reactions are catalytic, they increase both forward and reverse reaction rates. RNA degradation through Fe^2+^ cleavage should be weighed against potential RNA polymerization and the benefits of increased catalytic activity. The same dualism exists with Mg^2+^, but our work suggests higher stakes with Fe^2+^. At the extremes, without M^2+^, RNA cannot form complex folds and has few avenues for catalytic or functional activity while with excessive M^2+^ RNA is degraded. There theoretically exists some point of balance wherein M^2+^ is beneficially utilized with some frequency of disabling cleavage. Given the increased potency of cleavage with Fe^2+^ relative to M^2+^, this balancing point may be at a lower concentration of Fe^2+^ than Mg^2+^. However, enhanced cofactor characteristics of Fe^2+^ may allow RNA to access more functions using less metal. On early Earth, heightened RNA cleavage in the presence of Fe^2+^ if balanced by a similar rate of RNA resupply would allow functional space to be explored in short time. RNAs would be selected that could cooperate with or tolerate a potent metal. Fe^2+^ may have been a force for accelerated RNA evolution on early Earth.

### Fe^2+^ associates with rRNA in vivo

Exchange of non-native metals for native metals is well-known during purification of proteins (51). We observe analogous phenomena with rRNA. Fe^2+^ can exchange with Mg^2+^ (and vice versa) during purification of ribosomes. Ribosomes purified in either Fe^2+^ or Mg^2+^ associate with 500-1000 M^2+^ ions that match the type of ion in the purification buffers. Our data support the tight association and lack of exchange of around 9 M^2+^ per ribosome. This subset of M^2+^ do not exchange during purification. The number of non-exchangeable M^2+^ closely matches the number of M^2+^ identified previously as a special class of deeply buried and highly coordinated M^2+^ in dinuclear microclusters (M^2+^-μc’s) (16). Mg^2+^ ions in M^2+^-μc’s are directly chelated by multiple phosphate oxygens of the rRNA backbone and are substantially dehydrated. M^2+^-μc’s within the LSU provide a framework for the ribosome’s peptidyl transferase center, the site of protein synthesis in the ribosome, suggesting an essential and ancient role for M^2+^-μc’s in the ribosome. There are four dinuclear M^2+^-μc’s in the LSU and one in the SSU, accounting for 10 M^2+^ (16). Displacement of these M^2+^ would require large-scale changes in ribosomal conformation. In sum, there are ten M^2+^ per ribosome that are expected to be refractory to exchange. We hypothesize that this subset M^2+^ are contained in M^2+^ −μc’s, which can be occupied by either Mg^2+^ or Fe^2+^ (17), depending on growth conditions.

We also hypothesize that ribosomes harvested from aerobic cells have low Fe^2+^/Mg^2+^ ratios because of low intracellular Fe^2+^ availability and lability. This hypothesis is supported by our observation that the number of slow exchanging Fe^2+^ per ribosome from aerobic cells is near the baseline of our measurements. It appears that ribosomes harvested from pre-GOE conditions have high Fe^2+^/Mg^2+^ ratios because of high intracellular Fe^2+^ availability and lability, as indicated by the close match in the number of slowly exchanging Fe^2+^ per ribosome and the number of available M^2+^ sites in ribosomal M^2+^-μc’s. In these experiments we detect only the Fe^2+^ ions that do not exchange during purification.

### Anoxic Fe^2+^ degrades rRNA within ribosomes

rRNA from all four growth conditions showed partial hydrolysis when ribosomes were purified in anoxic Fe^2+^. It appears that Fe^2+^ can mediate rRNA degradation by an in-line mechanism during ribosomal purification in anoxic Fe^2+^. Less rRNA cleavage was observed in ribosomes purified with Mg^2+^, which contain orders of magnitude lower Fe^2+^.

### Summary

Here we have shown for the first time that bacteria grown in pre-GOE conditions contain functional ribosomes with tightly bound Fe atoms. The ~10 ribosomal Fe ions in ribosomes grown anoxically with high Fe^2+^ are likely deeply buried and specifically bound to rRNA. Depending on intracellular Fe lability, ribosomes may have higher Fe content *in vivo* given the high capacity for the ribosome to substitute ~600 loosely bound Mg^2+^ ions for Fe^2+^. Furthermore, direct association of the rRNA with Fe atoms results in a fast rate of in-line cleavage. 1 mM Fe^2+^ gives a ~10 times higher k_obs_ than does 25 mM Mg^2+^ so that the assumed per molar metal rate constant is hundreds of times greater with Fe^2+^ than with Mg^2+^. This highlights a potential role of protection from in-line cleavage for rProteins and suggests that Fe^2+^ may drive rapid cycling of RNA between monomers and polymers. Our results support a model in which alternate M^2+^ ions, namely Fe^2+^, participated in the origin and early evolution of life: first in abiotic proto-biochemical systems, through potentially rapid rounds of formation and breakdown of RNA structures, and then within early cellular life up until the GOE (65). Our study also expands the role of Fe^2+^ in modern biochemistry by showing that extant life retains the ability to incorporate Fe into ribosomes. We surmise that extant organisms under certain environmental and cellular states may use Fe^2+^ as a ribosomal cofactor. In addition, obligate anaerobic organisms that have spent the entirety of their evolutionary history in permanently anoxic environments may still use abundant Fe^2+^ in their ribosomes *in vivo*.

## Acknowledgments

We thank Corinna Tuckey (New England BioLabs), Eric B. O’Neill, and Drs. Anton Petrov, Roger M. Wartell, Thomas Tullius, and Ada Yonath for helpful discussions.

## References

1. Bernier, C.R., Petrov, A.S., Kovacs, N.A., Penev, P.I. and Williams, L.D. (2018) Translation: The Universal Structural Core of Life. Mol. Biol. Evol., 35, 2065–2076.

2. Melnikov, S., Ben-Shem, A., Garreau de Loubresse, N., Jenner, L., Yusupova, G. and Yusupov, M. (2012) One core, two shells: bacterial and eukaryotic ribosomes. Nat. Struct. Mol. Biol., 19, 560–567.

3. Woese, C.R. (2001) Translation: in retrospect and prospect. RNA, 7, 1055–1067.

4. Noller, H.F., Kop, J., Wheaton, V., Brosius, J., Gutell, R.R., Kopylov, A.M., Dohme, F., Herr, W., Stahl, D.A. and Gupta, R. (1981) Secondary structure model for 23S ribosomal RNA. Nucleic Acids Res., 9, 6167–6189.

5. Petrov, A.S., Bernier, C.R., Hsiao, C., Norris, A.M., Kovacs, N.A., Waterbury, C.C., Stepanov, V.G., Harvey, S.C., Fox, G.E., Wartell, R.M. et al. (2014) Evolution of the ribosome at atomic resolution. Proc. Natl. Acad. Sci. USA, 111, 10251–10256.

6. Bokov, K. and Steinberg, S.V. (2009) A hierarchical model for evolution of 23S ribosomal RNA. Nature, 457, 977–980.

7. Kovacs, N.A., Petrov, A.S., Lanier, K.A. and Williams, L.D. (2017) Frozen in Time: The History of Proteins. Mol. Biol. Evol., 34, 1252–1260.

8. Agmon, I., Bashan, A. and Yonath, A. (2006) On ribosome conservation and evolution. Isr. J. Ecol. Evol., 52, 359–374.

9. Klein, D.J., Moore, P.B. and Steitz, T.A. (2004) The contribution of metal ions to the structural stability of the large ribosomal subunit. RNA, 10, 1366–1379.

10. Anbar, A.D. (2008) Elements and evolution. Science, 322, 1481–1483.

11. Hazen, R.M. and Ferry, J.M. (2010) Mineral evolution: Mineralogy in the fourth dimension. Elements, 6, 9–12.

12. Holland, H.D. (2006) The oxygenation of the atmosphere and oceans. Philos. Trans. R. Soc. London, Ser. B, 361, 903–915.

13. Klein, C. (2005) Some Precambrian banded iron-formations (BIFs) from around the world: Their age, geologic setting, mineralogy, metamorphism, geochemistry, and origins. Am. Mineral., 90, 1473–1499.

14. Holland, H.D. (1973) The oceans; a possible source of iron in iron-formations. Econ. Geol., 68, 1169–1172.

15. Bowman, J.C., Lenz, T.K., Hud, N.V. and Williams, L.D. (2012) Cations in charge: magnesium ions in RNA folding and catalysis. Curr. Opin. Struct. Biol., 22, 262–272.

16. Hsiao, C. and Williams, L.D. (2009) A recurrent magnesium-binding motif provides a framework for the ribosomal peptidyl transferase center. Nucleic Acids Res., 37, 3134–3142.

17. Lin, S.Y., Wang, Y.C. and Hsiao, C. (2019) Prebiotic Iron Originates the Peptidyl Transfer Origin. Mol. Biol. Evol., 36, 999–1007.

18. Schuwirth, B.S., Borovinskaya, M.A., Hau, C.W., Zhang, W., Vila-Sanjurjo, A., Holton, J.M. and Cate, J.H.D. (2005) Structures of the Bacterial Ribosome at 3.5 Å Resolution. Science, 310, 827–834.

19. Selmer, M., Dunham, C.M., Murphy, F.V., Weixlbaumer, A., Petry, S., Kelley, A.C., Weir, J.R. and Ramakrishnan, V. (2006) Structure of the 70S ribosome complexed with mRNA and tRNA. Science, 313, 1935–1942.

20. Demeshkina, N., Jenner, L., Westhof, E., Yusupov, M. and Yusupova, G. (2012) A new understanding of the decoding principle on the ribosome. Nature, 484, 256–259.

21. Petrov, A.S., Bernier, C.R., Hsiao, C., Okafor, C.D., Tannenbaum, E., Stern, J., Gaucher, E., Schneider, D., Hud, N.V., Harvey, S.C. et al. (2012) RNA-magnesium-protein interactions in large ribosomal subunit. J. Phys. Chem. B, 116, 8113–8120.

22. Forconi, M. and Herschlag, D. (2009) Metal ion-based RNA cleavage as a structural probe. Methods Enzymol., 468, 91–106.

23. Soukup, G.A. and Breaker, R.R. (1999) Relationship between internucleotide linkage geometry and the stability of RNA. RNA, 5, 1308–1325.

24. Winkler, W., Nahvi, A. and Breaker, R.R. (2002) Thiamine derivatives bind messenger RNAs directly to regulate bacterial gene expression. Nature, 419, 952–956.

25. Winkler, W.C., Nahvi, A., Roth, A., Collins, J.A. and Breaker, R.R. (2004) Control of gene expression by a natural metabolite-responsive ribozyme. Nature, 428, 281.

26. Dorner, S. and Barta, A. (1999) Probing ribosome structure by europium-induced RNA cleavage. Biol. Chem., 380, 243–251.

27. Winter, D., Polacek, N., Halama, I., Streicher, B. and Barta, A. (1997) Lead-catalysed specific cleavage of ribosomal RNAs. Nucleic Acids Res., 25, 1817–1824.

28. Pyle, A.M. (2002) Metal ions in the structure and function of RNA. J. Biol. Inorg. Chem., 7, 679–690.

29. Pan, T. and Uhlenbeck, O.C. (1992) In vitro selection of RNAs that undergo autolytic cleavage with lead (2+). Biochemistry, 31, 3887–3895.

30. Winterbourn, C.C. (1995) Toxicity of iron and hydrogen peroxide: the Fenton reaction. Toxicol. Lett., 82, 969–974.

31. Fischbacher, A., von Sonntag, C. and Schmidt, T.C. (2017) Hydroxyl radical yields in the Fenton process under various pH, ligand concentrations and hydrogen peroxide/Fe(II) ratios. Chemosphere, 182, 738–744.

32. Dixon, W.J., Hayes, J.J., Levin, J.R., Weidner, M.F., Dombroski, B.A. and Tullius, T.D. (1991) Hydroxyl radical footprinting. Methods Enzymol., 208, 380–413.

33. Tullius, T.D. (1996) In Suslick, K. (ed.), Comprehensive Supramolecular Chemistry. Elsevier, Tarrytown, NY, Vol. 5, pp. 317–343.

34. Celander, D.W. and Cech, T.R. (1991) Visualizing the higher order folding of a catalytic RNA molecule. Science, 251, 401–407.

35. Li, Z., Wu, J. and Deleo, C.J. (2006) RNA damage and surveillance under oxidative stress. IUBMB Life, 58, 581–588.

36. Shcherbik, N. and Pestov, D.G. (2019) The Impact of Oxidative Stress on Ribosomes: From Injury to Regulation. Cells, 8, 1379.

37. Bray, M.S., Lenz, T.K., Haynes, J.W., Bowman, J.C., Petrov, A.S., Reddi, A.R., Hud, N.V., Williams, L.D. and Glass, J.B. (2018) Multiple prebiotic metals mediate translation. Proc. Natl. Acad. Sci. USA, 115, 12164–12169.

38. Li, Y. and Breaker, R.R. (1999) Kinetics of RNA Degradation by Specific Base Catalysis of Transesterification Involving the 2’-Hydroxyl Group. J. Am. Chem. Soc., 121, 5364–5372.

39. Kuusela, S. and Lönnberg, H. (1993) Metal ions that promote the hydrolysis of nucleoside phosphoesters do not enhance intramolecular phosphate migration. J. Phys. Org., 6, 347–356.

40. Hsiao, C., Lenz, T.K., Peters, J.K., Fang, P.-Y., Schneider, D.M., Anderson, E.J., Preeprem, T., Bowman, J.C., O’Neill, E.B. and Lie, L. (2013) Molecular paleontology: a biochemical model of the ancestral ribosome. Nucleic Acids Res., 41, 3373–3385.

41. Riemer, J., Hoepken, H.H., Czerwinska, H., Robinson, S.R. and Dringen, R. (2004) Colorimetric ferrozine-based assay for the quantitation of iron in cultured cells. Anal. Biochem., 331, 370–375.

42. Maguire, B.A., Wondrack, L.M., Contillo, L.G. and Xu, Z. (2008) A novel chromatography system to isolate active ribosomes from pathogenic bacteria. RNA, 14, 188–195.

43. Shimizu, Y., Inoue, A., Tomari, Y., Suzuki, T., Yokogawa, T., Nishikawa, K. and Ueda, T. (2001) Cell-free translation reconstituted with purified components. Nature Biotechnol., 19, 751–755.

44. Rappsilber, J., Mann, M. and Ishihama, Y. (2007) Protocol for micro-purification, enrichment, pre-fractionation and storage of peptides for proteomics using StageTips. Nat Protoc, 2, 1896–1906.

45. Cox, J. and Mann, M. (2008) MaxQuant enables high peptide identification rates, individualized p.p.b.-range mass accuracies and proteome-wide protein quantification. Nat. Biotechnol., 26, 1367–1372.

46. Cox, J., Neuhauser, N., Michalski, A., Scheltema, R.A., Olsen, J.V. and Mann, M. (2011) Andromeda: a peptide search engine integrated into the MaxQuant environment. J. Proteome. Res., 10, 1794–1805.

47. Kurylo, C.M., Alexander, N., Dass, R.A., Parks, M.M., Altman, R.A., Vincent, C.T., Mason, C.E. and Blanchard, S.C. (2016) Genome Sequence and Analysis of *Escherichia coli* MRE600, a Colicinogenic, Nonmotile Strain that Lacks RNase I and the Type I Methyltransferase, EcoKI. Genome Biol. Evol., 8, 742–752.

48. Tyanova, S., Temu, T., Sinitcyn, P., Carlson, A., Hein, M.Y., Geiger, T., Mann, M. and Cox, J. (2016) The Perseus computational platform for comprehensive analysis of (prote)omics data. Nat. Methods, 13, 731–740.

49. Tullius, T.D., Dombroski, B.A., Churchill, M.E. and Kam, L. (1987) Hydroxyl radical footprinting: A high-resolution method for mapping protein-DNA contacts. Methods Enzymol., 155, 537–558.

50. Pogozelski, W.K. and Tullius, T.D. (1998) Oxidative Strand Scission of Nucleic Acids: Routes Initiated by Hydrogen Abstraction from the Sugar Moiety. Chem. Rev., 98, 1089–1108.

51. Handing, K.B., Niedzialkowska, E., Shabalin, I.G., Kuhn, M.L., Zheng, H. and Minor, W. (2018) Characterizing metal-binding sites in proteins with X-ray crystallography. Nat. Protoc., 13, 1062.

52. Rivera, M.C., Maguire, B. and Lake, J.A. (2015) Isolation of Ribosomes and Polysomes. Cold Spring Harb. Protoc., 2015, pdb.prot081331.

53. Kuusela, S., Azhayev, A., Guzaev, A. and Lönnberg, H. (1995) The effect of the 3’-terminal monophosphate group on the metal-ion-promoted hydrolysis of the phosphodiester bonds of short oligonucleotides. J. Chem. Soc. Perk. Trans. 2, 1197–1202.

54. Kuusela, S. and Lönnberg, H. (1996) Zn^2+^-promoted hydrolysis of 3’,5’-dinucleoside monophosphates and polyribonucleotides. The effect of nearest neighbours on the cleavage of phosphodiester bonds. Nucleosides Nucleotides Nucl. Acids, 15, 1669–1678.

55. Kuusela, S. and Lönnberg, H. (1994) Metal-ion-promoted hydrolysis of polyuridylic acid. J. Chem. Soc. Perk. Trans. 2, 2301–2306.

56. Ikenaga, H. and Inoue, Y. (1974) Metal (II) ion catalyzed transphosphorylation of four homodinucleotides and five pairs of dinucleotide sequence isomers. Biochemistry, 13, 577–582.

57. Breslow, R. and Huang, D.-L. (1991) Effects of metal ions, including Mg^2+^ and lanthanides, on the cleavage of ribonucleotides and RNA model compounds. Proc. Natl. Acad. Sci. USA, 88, 4080–4083.

58. Honda, K., Smith, M.A., Zhu, X., Baus, D., Merrick, W.C., Tartakoff, A.M., Hattier, T., Harris, P.L., Siedlak, S.L., Fujioka, H. et al. (2005) Ribosomal RNA in Alzheimer disease is oxidized by bound redox-active iron. J Biol. Chem., 280, 20978–20986.

59. Zinskie, J.A., Ghosh, A., Trainor, B.M., Shedlovskiy, D., Pestov, D.G. and Shcherbik, N. (2018) Iron-dependent cleavage of ribosomal RNA during oxidative stress in the yeast *Saccharomyces cerevisiae*. J Biol. Chem., 293, 14237–14248.

60. Athavale, S.S., Petrov, A.S., Hsiao, C., Watkins, D., Prickett, C.D., Gossett, J.J., Lie, L., Bowman, J.C., O’Neill, E. and Bernier, C.R. (2012) RNA folding and catalysis mediated by iron (II). PLoS One, 7, e38024.

61. Moon, W.J. and Liu, J. (2020) Replacing Mg^2+^ by Fe^2+^ for RNA-Cleaving DNAzymes. ChemBioChem, 21, 401–407.

62. Okafor, C.D., Lanier, K.A., Petrov, A.S., Athavale, S.S., Bowman, J.C., Hud, N.V. and Williams, L.D. (2017) Iron mediates catalysis of nucleic acid processing enzymes: Support for Fe(II) as a cofactor before the Great Oxidation Event. Nucleic Acids Res., 45, 3634–3642.

63. Hsiao, C., Chou, I.-C., Okafor, C.D., Bowman, J.C., O’neill, E.B., Athavale, S.S., Petrov, A.S., Hud, N.V., Wartell, R.M., Harvey, S.C. et al. (2013) RNA with iron (II) as a cofactor catalyses electron transfer. Nat. Chem., 5, 525–528.

64. Ward, W.L., Plakos, K. and DeRose, V.J. (2014) Nucleic acid catalysis: metals, nucleobases, and other cofactors. Chemical Reviews, 114, 4318–4342.

65. Okafor, C.D., Bowman, J.C., Hud, N.V., Glass, J.B. and Williams, L.D. (2018), Prebiotic Chemistry and Chemical Evolution of Nucleic Acids. Springer, pp. 227–243.

