## Supplemental Materials for "Cutting in-line with iron: ribosomal function and non-oxidative RNA cleavage"

**Supplemental Information:**

Supplemental Tables S1-S3

Supplemental Figures S1-S7

**SUPPLEMENTAL TABLES**

**Table S1.** **Proteins differentially expressed in ribosomes purified from aerobic vs anaerobic cells.** Information on metal cofactors was taken from the UniProt entries for each protein.

| **Condition** | **Protein more abundant in** | **Protein name** | **UniProt ID** | **Fold more abundant** | **p-value** | **Metal cofactor** |
| --- | --- | --- | --- | --- | --- | --- |
| 1 mM added Fe^2+^ | Anaerobic  vs.  Aerobic | L-threonine dehydratase catabolic TdcB | P0AGF6 | 211.8 | 0.02 | N.A. |
|  |  | Hydrogenase maturation factor HypB | P0AAN3 | 12.1 | 0.03 | N.A. |
|  |  | Glycerol dehydrogenase | P0A9S5 | 11.4 | 0.01 | Zn^2+^ |
|  |  | Phosphoribosylaminoimidazole-succinocarboxamide synthase | P0A7D7 | 6.0 | 0.02 | N.A. |
|  |  | Aldehyde-alcohol dehydrogenase | P0A9Q7 | 6.0 | 0.03 | Fe^2+^ |
|  |  | Uncharacterized protein YjjI | P37342 | 5.8 | 0.02 | N.A. |
|  |  | Probable acrylyl-CoA reductase AcuI | P26646 | 5.5 | 0.01 | N.A. |
|  |  | Bacterial non-heme ferritin^a^ | P0A998 | 5.1 | 0.03 | N.A. |
|  |  | Aspartate ammonia-lyase | P0AC38 | 4.1 | 0.01 | N.A. |
|  |  | Fumarate reductase flavoprotein subunit | P00363 | 3.7 | 0.05 | N.A. |
|  |  | UvrABC system protein B | P0A8F8 | 3.6 | 0.01 | N.A. |
|  |  | Formate acetyltransferase 1 | P09373 | 3.2 | 0.01 | N.A. |
|  |  | UPF0227 protein YcfP | P0A8E1 | 3.1 | 0.02 | N.A. |
|  |  | Phosphoenolpyruvate-protein phosphotransferase | P08839 | 2.6 | 0.03 | Mg^2+^ |
|  |  | Ubiquinone/menaquinone biosynthesis C-methyltransferase UbiE | P0A887 | 2.4 | 0.02 | N.A. |
|  |  | Glycogen phosphorylase | P0AC86 | 2.2 | 0.01 | N.A. |
|  |  | Maltodextrin phosphorylase | P00490 | 2.1 | 0.04 | N.A. |
|  |  | Thymidine phosphorylase | P07650 | 2.1 | 0.01 | N.A. |
|  |  | Protein-export protein SecB | P0AG86 | 2.0 | 0.01 | N.A. |
|  |  | Integration host factor subunit beta | P0A6Y1 | 2.0 | 0.04 | N.A. |
|  |  | Cold shock-like protein CspE | P0A972 | 2.0 | 0.03 | N.A. |
|  |  | Fatty acid metabolism regulator protein | P0A8V6 | 1.9 | 0.02 | N.A. |
|  | Aerobic  vs.  Anaerobic | Lipoyl synthase | P60716 | 2.2 | 0.04 | [4Fe-4S] |
|  |  | Translation initiation factor IF-3^b^ | P0A707 | 2.2 | 0.00 | N.A. |
|  |  | Ribosomal protein S12 methylthiotransferase RimO^b^ | P0AEI4 | 2.3 | 0.02 | [4Fe-4S] |
|  |  | Dihydrolipoyllysine-residue acetyltransferase component of pyruvate dehydrogenase complex | P06959 | 2.4 | 0.01 | N.A. |
|  |  | Ribonucleotide monophosphatase NagD | P0AF24 | 2.4 | 0.01 | Mg^2+^ |
|  |  | Ribosomal RNA large subunit methyltransferase K/L^b^ | P75864 | 2.4 | 0.04 | N.A. |
|  |  | 2-oxoglutarate dehydrogenase E1 component | P0AFG3 | 4.7 | 0.01 | N.A. |
|  |  | Alkyl hydroperoxide reductase C | P0AE08 | 5.4 | 0.05 | N.A. |
|  |  | Pyruvate/proton symporter BtsT | P39396 | 9.1 | 0.02 | N.A. |
|  |  | Bifunctional protein PutA | P09546 | 10.7 | 0.02 | N.A. |
| No added Fe^2+^ | Anaerobic  vs.  Aerobic | L-threonine dehydratase catabolic TdcB | P0AGF6 | 137.5 | 0.00 | N.A. |
|  |  | Uncharacterized protein YfdQ | P76513 | 38.0 | 0.02 | N.A. |
|  |  | Hydrogenase-2 large chain | P0ACE0 | 31.5 | 0.01 | Ni^2+^ |
|  |  | Fumarate reductase iron-sulfur subunit | P0AC47 | 17.5 | 0.01 | [2Fe-2S],  [3Fe-4S],  [4Fe-4S] |
|  |  | Glycerol dehydrogenase | P0A9S5 | 16.9 | 0.01 | Zn^2+^ |
|  |  | Bacterial non-heme ferritin^a^ | P0A998 | 14.9 | 0.00 | N.A. |
|  |  | Hydrogenase maturation factor HypB | P0AAN3 | 11.6 | 0.02 | N.A. |
|  |  | PFL-like enzyme TdcE | P42632 | 8.6 | 0.01 | N.A. |
|  |  | Aldehyde-alcohol dehydrogenase | P0A9Q7 | 7.2 | 0.01 | Fe^2+^ |
|  |  | Anaerobic ribonucleoside-triphosphate reductase | P28903 | 5.8 | 0.04 | N.A. |
|  |  | Aspartate ammonia-lyase | P0AC38 | 4.5 | 0.01 | N.A. |
|  |  | Uncharacterized protein YjjI | P37342 | 4.3 | 0.03 | N.A. |
|  |  | SCP2 domain-containing protein YhbT | P64599 | 4.1 | 0.01 | N.A. |
|  |  | 2,3-bisphosphoglycerate-independent phosphoglycerate mutase | P37689 | 3.9 | 0.00 | Mn^2+^ |
|  |  | 2-octaprenylphenol hydroxylase | P25535 | 3.7 | 0.05 | N.A. |
|  |  | Fructose-1,6-bisphosphatase 1 class 2 | P0A9C9 | 3.3 | 0.03 | Mn^2+^ |
|  |  | Stationary-phase-induced ribosome-associated protein^b^ | P68191 | 3.3 | 0.00 | N.A. |
|  |  | Evolved beta-galactosidase subunit alpha | P06864 | 3.2 | 0.04 | N.A. |
|  |  | Ribonuclease T | P30014 | 2.8 | 0.03 | Mg^2+^ |
|  |  | UPF0227 protein YcfP | P0A8E1 | 2.5 | 0.03 | N.A. |
|  |  | UPF0313 protein YgiQ | Q46861 | 2.4 | 0.01 | [4Fe-4S] |
|  |  | Methionine--tRNA ligase | P00959 | 2.4 | 0.02 | Zn^2+^ |
|  |  | Polyphosphate kinase | P0A7B1 | 2.4 | 0.03 | Mg^2+^ |
|  |  | RecBCD enzyme subunit RecC | P07648 | 2.3 | 0.02 | N.A. |
|  |  | ADP-heptose--LPS heptosyltransferase 2 | P37692 | 2.2 | 0.03 | N.A. |
|  | Aerobic  vs.  Anaerobic | RNA-binding protein YhbY | P0AGK4 | 2.2 | 0.01 | N.A. |
|  |  | Chaperone protein HscA | P0A6Z1 | 2.3 | 0.00 | N.A. |
|  |  | Cytochrome bd-I ubiquinol oxidase subunit 2 | P0ABK2 | 2.3 | 0.03 | heme |
|  |  | 5'-methylthioadenosine/S-adenosylhomocysteine nucleosidase | P0AF12 | 2.3 | 0.02 | N.A. |
|  |  | Ribosomal silencing factor RsfS^b^ | P0AAT6 | 2.4 | 0.00 | N.A. |
|  |  | DNA topoisomerase 4 subunit B | P20083 | 2.7 | 0.01 | Mg^2+^ |
|  |  | Lipoyl synthase | P60716 | 3.0 | 0.01 | [4Fe-4S] |
|  |  | Transcriptional regulatory protein GlrR | P0AFU4 | 3.2 | 0.00 | N.A. |
|  |  | Modulator of FtsH protease HflC | P0ABC3 | 3.5 | 0.01 | N.A. |
|  |  | Ribosomal RNA small subunit methyltransferase A^b^ | P06992 | 3.7 | 0.01 | N.A. |
|  |  | Pyruvate/proton symporter BtsT | P39396 | 5.8 | 0.04 | N.A. |
|  |  | Enterobactin synthase component B^a^ | P0ADI4 | 8.0 | 0.01 | Mg^2+^ |
|  |  | Respiratory nitrate reductase 1 alpha chain | P09152 | 37.3 | 0.01 | [4Fe-4S]  Mo-bis-MGD |

^a^Protein is involved in bacterial iron homeostasis; ^b^Protein is involved in translation

**Table S2. Proteins differentially expressed in ribosomes purified from cells grown with or without 1 mM added Fe^2+^.** Information on metal cofactors was taken from the UniProt entries for each protein.

| **Condition** | **Protein more abundant in** | **Protein name** | **UniProt ID** | **Fold more abundant** | **p-value** | **Metal**  **cofactor** |
| --- | --- | --- | --- | --- | --- | --- |
| Anaerobic | 1 mM added Fe  vs.  without added Fe | Transcriptional regulatory protein OmpR | P0AA16 | 2.0 | 0.01 | N.A. |
|  |  | UvrABC system protein B | P0A8F8 | 2.1 | 0.03 | N.A. |
|  |  | 5'-methylthioadenosine/S-adenosylhomocysteine nucleosidase | P0AF12 | 2.3 | 0.04 | N.A. |
|  |  | Large ribosomal RNA subunit accumulation protein YceD^b^ | P0AB28 | 2.3 | 0.01 | N.A. |
|  |  | ATP-dependent dethiobiotin synthetase BioD 2 | P0A6E9 | 2.7 | 0.04 | Mg^2+^ |
|  |  | Phosphoribosylaminoimidazole-succinocarboxamide synthase | P0A7D7 | 2.7 | 0.01 | N.A. |
|  |  | Phosphoserine phosphatase | P0AGB0 | 4.1 | 0.02 | Mg^2+^ |
|  | Without added Fe  vs. 1 mM added Fe | 4-alpha-glucanotransferase | P15977 | 2.2 | 0.03 | N.A. |
| Aerobic | 1 mM added Fe  vs.  without added Fe | Probable 4-deoxy-4-formamido-L-arabinose-phosphoundecaprenol deformylase ArnD | P76472 | 2.0 | 0.01 | N.A. |
|  |  | GDP-mannose pyrophosphatase NudK | P37128 | 2.0 | 0.03 | Mg^2+^ |
|  |  | Uncharacterized protein YfdQ | P76513 | 2.1 | 0.02 | N.A. |
|  |  | Uncharacterized protein YciO | P0AFR4 | 2.1 | 0.00 | N.A. |
|  |  | ABC transporter ATP-binding protein ModF | P31060 | 3.1 | 0.04 | N.A. |
|  | Without added Fe  vs. 1 mM added Fe | Enterobactin synthase component B^a^ | P0ADI4 | 8.9 | 0.03 | Mg^2+^ |
|  |  | Fe(3+) dicitrate-binding periplasmic protein^a^ | P15028 | 7.7 | 0.02 | N.A. |

^a^Protein is involved in bacterial iron homeostasis

^b^Protein is involved in translation

| **Table S3: Reaction kinetics of in-line cleavage** | | | | | |
| --- | --- | --- | --- | --- | --- |
|  | Conditions | k_obs_ (10^-5^ sec^-1^) | No. cleavable phosphates | [M^2+^] (M) | k/phosphate (10^-5^ sec^-1^, no. phosphates^-1^, [M^2+^]^-1^) |
| This study | | | | | |
| Mg^2+^, 23S rRNA | pH 7.6, 37°C | 5 | 2905 | 0.025 | 0.07 |
| Mg^2+^, 16S rRNA | pH 7.6, 37°C | 3 | 1541 | 0.025 | 0.08 |
| Fe^2+^, 23S rRNA | pH 7.6, 37°C | 67 | 2905 | 0.001 | 23 |
| Fe^2+^, 16S rRNA | pH 7.6, 37°C | 25 | 1541 | 0.001 | 16 |
| Outside studies | | | | | |
| ^a^Mg^2+^, UpU | pH 5.6, 90°C | 0.01 | 1 | 0.005 | 2.1 |
| ^b^Mg^2+^, *S(ApG) | pH 9.5, 37°C | 0.18 | 1 | 0.005 | 37 |
| ^c^Mg^2+^, polyU | pH 5.6, 90°C | 0.32 | 300 | 0.005 | 0.21 |
| ^d^Zn^2+^, ApA | pH 5.1, 90°C | 0.64 | 1 | 0.01 | 64 |
| ^e^Zn^2+^, ApA | pH 7.0, 62.1°C | 0.06 | 1 | 0.001 | 56 |
| ^a^Zn^2+^, UpU | pH 5.6, 90°C | 0.43 | 1 | 0.005 | 85 |
| ^f^Zn^2+^, UpU | pH 7, 80°C | 0.04 | 1 | 0.001 | 41 |
| ^c^Zn^2+^, polyU | pH 5.6, 90°C | 0.43 | 300 | 0.005 | 0.28 |
| ^g^Zn^2+^, Up(Tp)_7_Tp | pH 5.5, 37°C | 0.50 | 1 | 0.01 | 50 |

^a^Kuusela & Lönnberg, 1993; ^b^Li & Breaker, 1999; ^c^Kuusela & Lönnberg, 1994; ^d^Kuusela & Lönnberg, 1996; ^e^Ikenaga & Inoue, 1974; ^f^Breslow & Huang, 1991, ^g^Kuusela et al., 1995.

*22 nucleotide DNA-RNA hybrid

**SUPPLEMENTAL FIGURES**

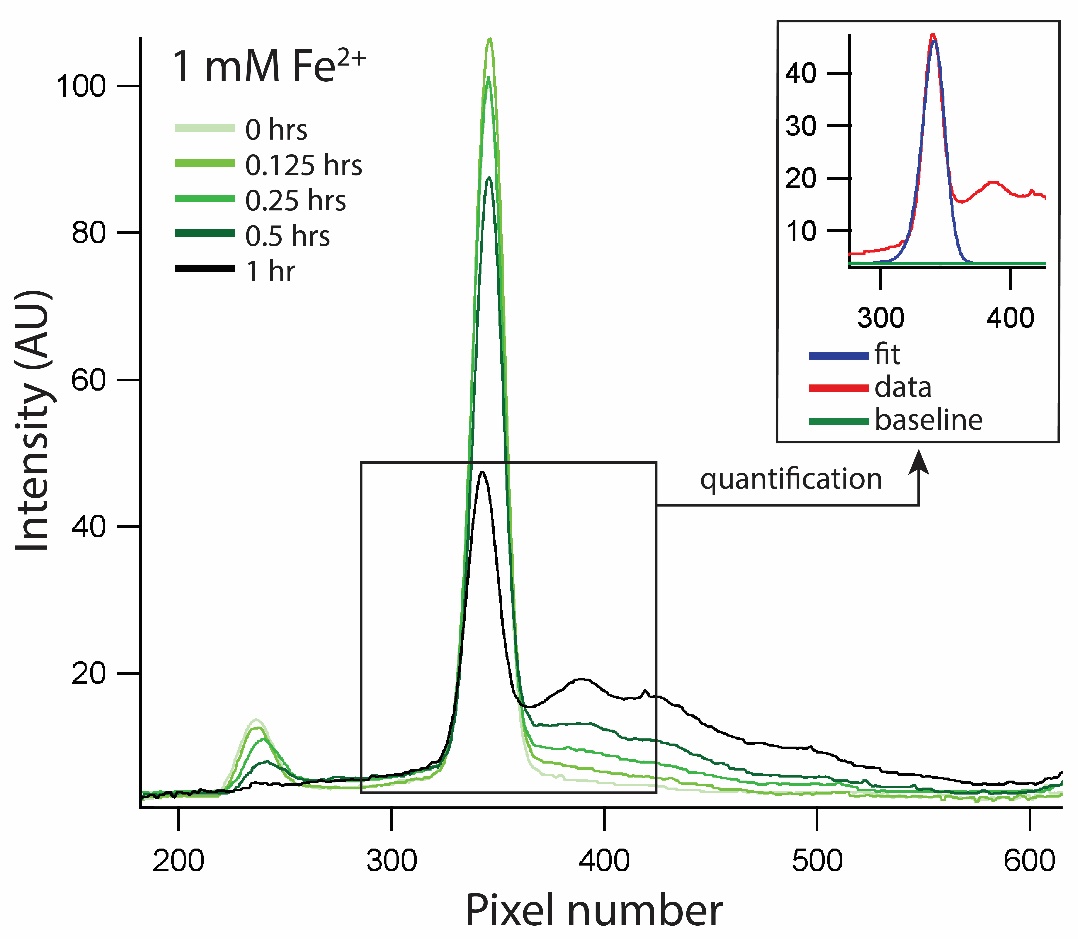

**Figure S1. Example quantification of a gel scan for rRNA reactions with 1 mM Fe^2+^**. Shown here is a typical example of lane profiles and peak quantification for rRNA in-line cleavage reactions from a gel scan. Peak fitting was done in Igor Pro 7. The areas of quantified peaks were compiled and plotted in Figure 1.

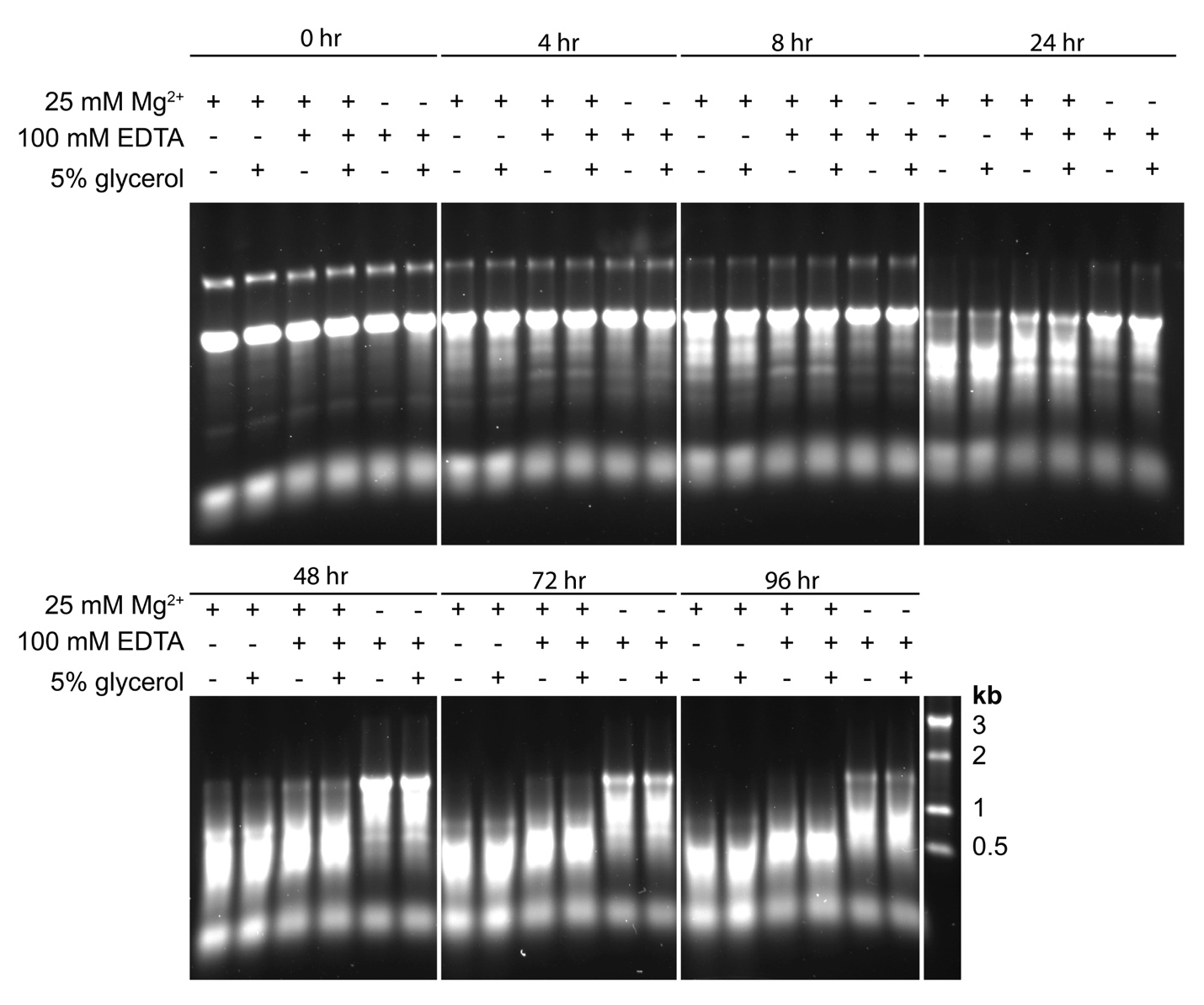

**Figure S2. 1% agarose gel showing 5% v/v glycerol does not inhibit Mg^2+^ in-line cleavage of naked rRNA at 37⁰C in air over the course of** **96 hours**. Chelation by 100 mM EDTA inhibits in-line cleavage, but there is no difference with and without 5% glycerol.

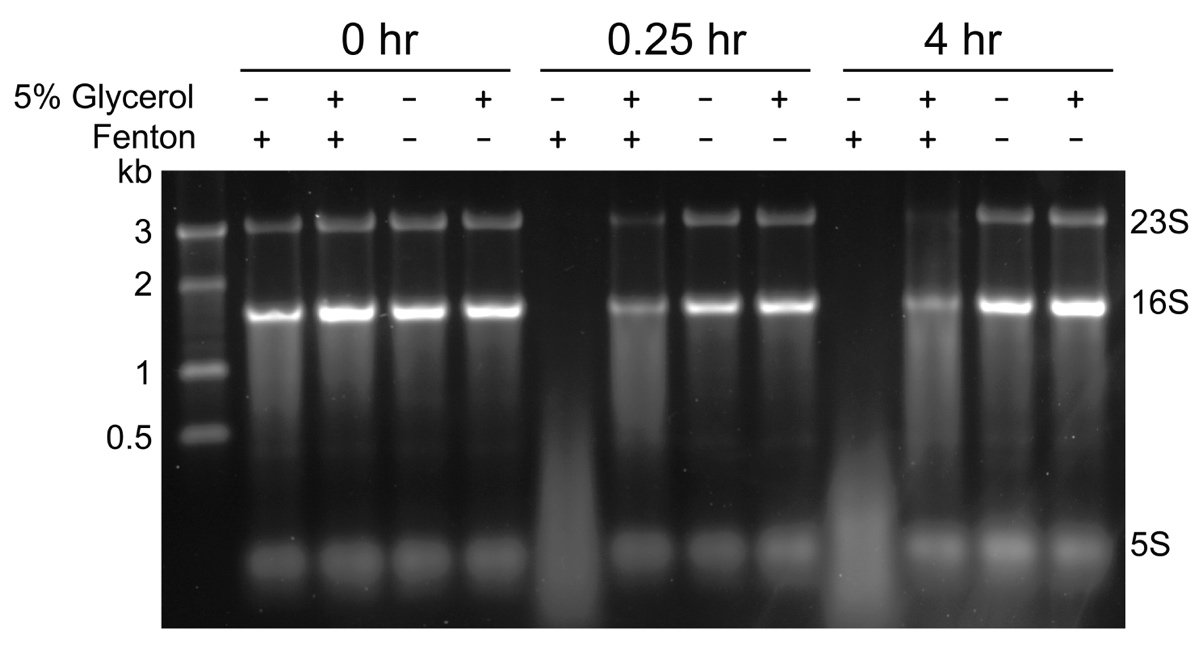

**Figure S3. 1% agarose gel showing 5% v/v glycerol inhibition of Fenton chemistry against naked rRNA at 37⁰C in air over the course of** **4 hours**. “+ Fenton” reactions contained 1 mM Fe^2+^, 0.3% H_2_O_2_, 10 mM ascorbic acid, and 10 mM EDTA. “- Fenton” reactions contained 10 mM EDTA.

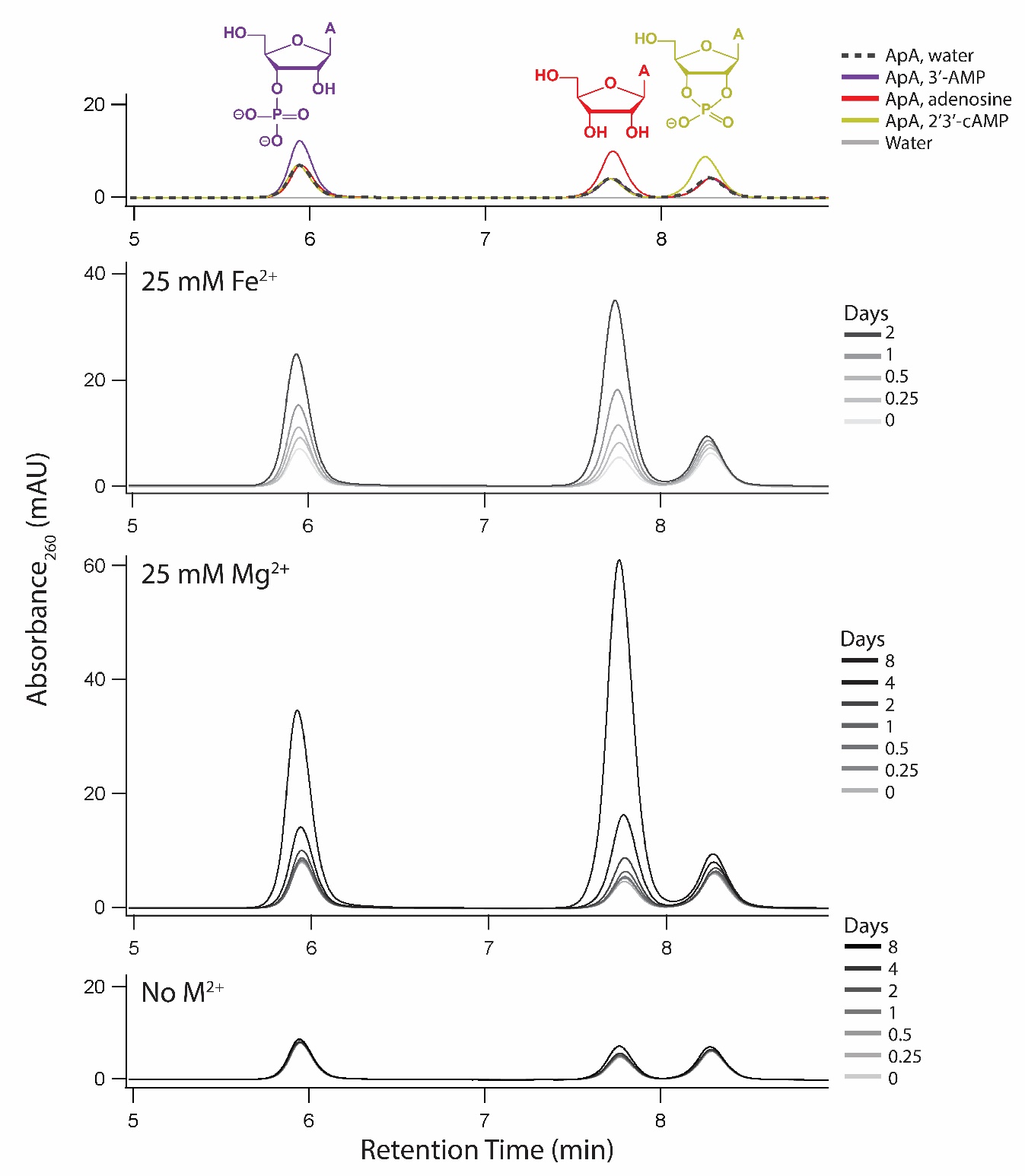
**Figure S4. C18 column HPLC chromatograms of ApA reactions showing accumulation of the same products with Fe^2+^ and Mg^2+^.** 3’-AMP, adenosine, and 2’,3’-cAMP standards (2.5 μM) were spiked into ApA solutions (0.5 mM ApA, 20 mM HEPES pH 7.6, 30 mM NaCl, 5% v/v glycerol) giving characteristic retention times for 3’-AMP, adenosine, and 2’,3’-cAMP. ApA solutions (0.5 mM ApA, 20 mM HEPES pH 7.6, 30 mM NaCl, 5% v/v glycerol) anoxically incubated at 37°C with either 25 mM Fe^2+^ out to 2 days or 25 mM Mg^2+^ out to 8 days accumulate 3’-AMP, adenosine, and 2’,3’-cAMP. 2’,3’-cAMP peaks build to a lesser extent due to cyclic phosphate hydrolysis to form 3’-AMP or 2’-AMP. There is little cleavage product formation in the no metal (M^2+^) control.

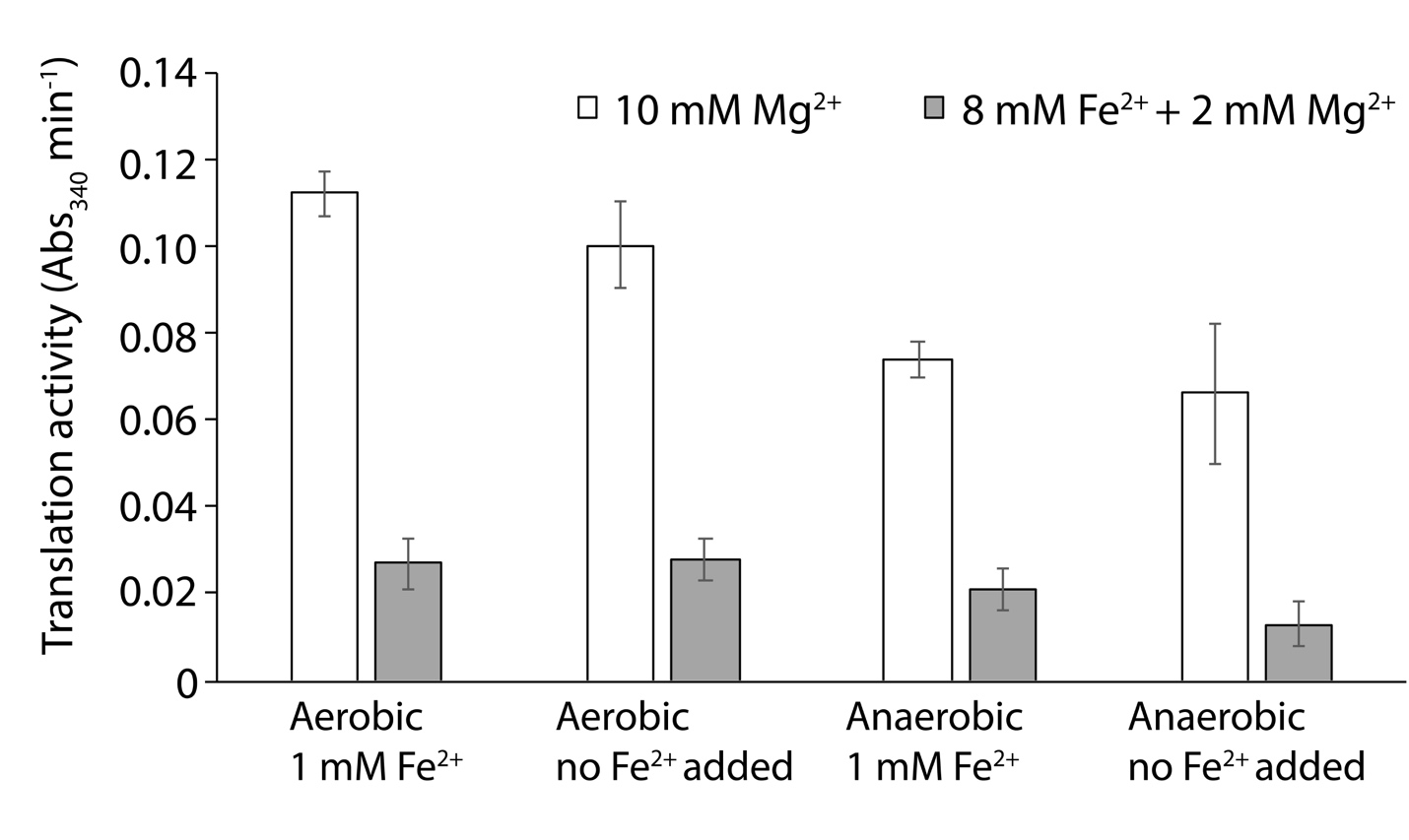

**Figure S5. *In vitro* translation activity of purified ribosomes.** Production of the protein dihydrofolate reductase (DHFR) from its mRNA was used to monitor translational activity. Protein synthesis was assayed by measuring the rate of NADPH oxidation at Abs_340_ by DHFR. Average values are reported ± standard error of the mean (n=4). All ribosomes were normalized to 9 mg mL^-1^ before adding to translation reactions.

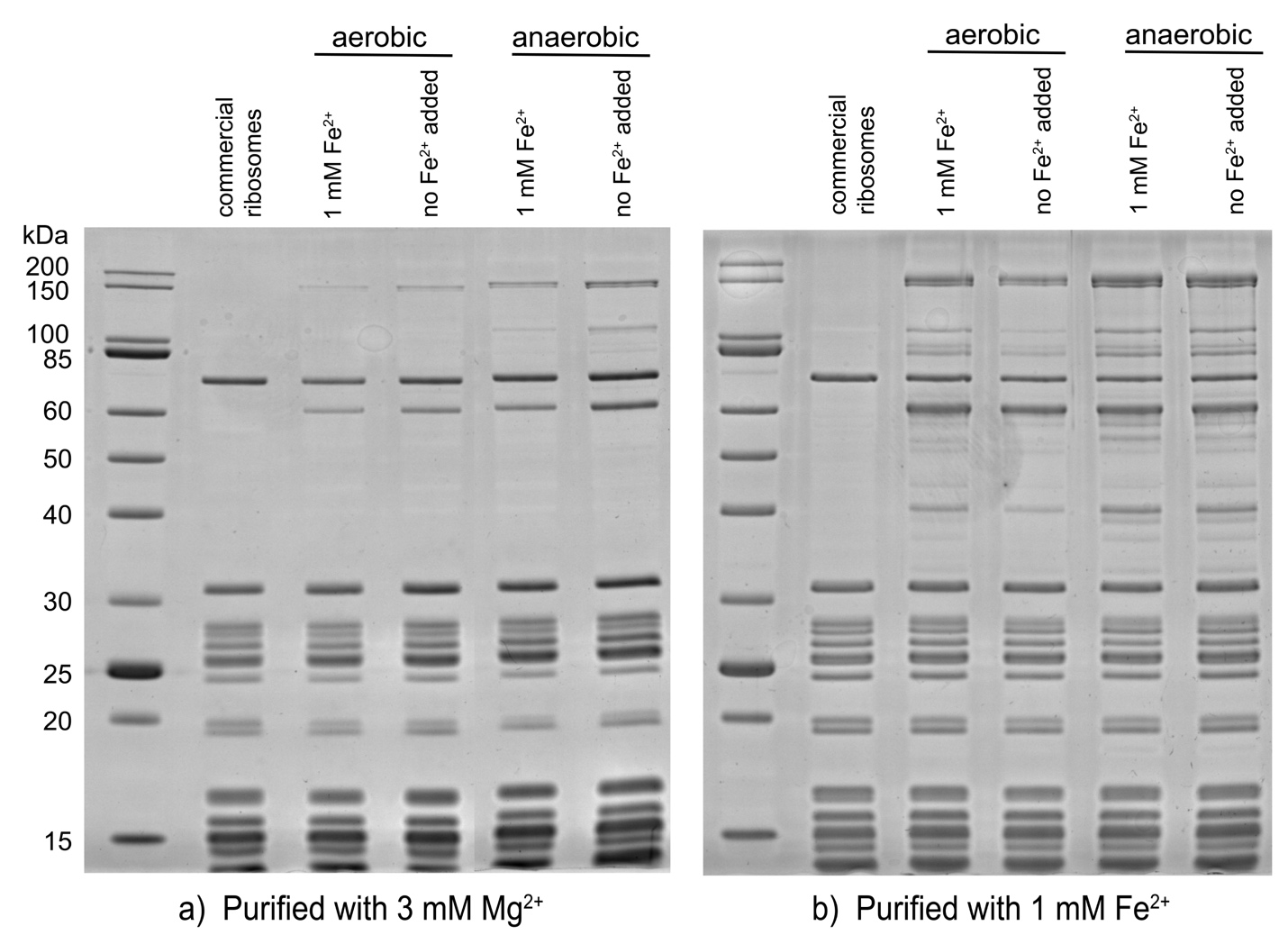

**Figure S6.** **12% SDS polyacrylamide gels for proteins from ribosomes purified in (a) 3 mM Mg^2+^ or (b) 1 mM Fe^2+^ compared to commercial ribosomes supplied by New England Biolabs.**

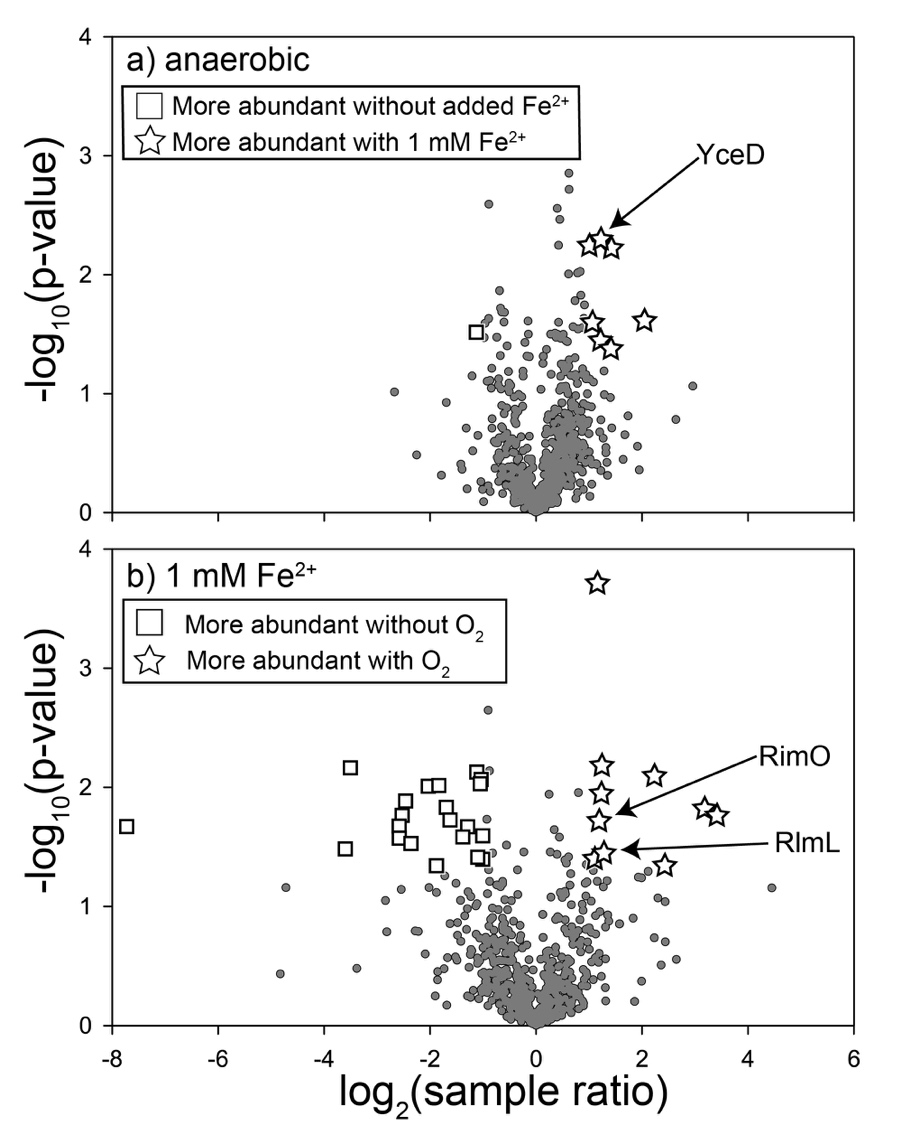

**Figure S7. Differential protein abundance between ribosomes purified from cells grown under four growth conditions.** Graphs display relative protein abundance in ribosome samples between two growth conditions. Black circles represent proteins not significantly more abundant in either sample. Gray rectangle and white stars represent proteins significantly more abundant in one of the samples. Proteins with a 2-fold or greater abundance in one sample versus another and a p-value less than or equal to 0.05 (n=3), were classified as significantly more abundant.
